# Accelerated spike-triggered non-negative matrix factorization reveals coordinated ganglion cell subunit mosaics in the primate retina

**DOI:** 10.1101/2024.04.22.590506

**Authors:** Sören J. Zapp, Mohammad H. Khani, Helene M. Schreyer, Shashwat Sridhar, Varsha Ramakrishna, Steffen Krüppel, Matthias Mietsch, Dario A. Protti, Dimokratis Karamanlis, Tim Gollisch

**Affiliations:** University Medical Center Göttingen, Department of Ophthalmology, Göttingen, Germany; Bernstein Center for Computational Neuroscience, Göttingen, Germany; International Max Planck Research School for Neurosciences, Göttingen, Germany; Collaborative Research Center CRC 1690 “Disease Mechanisms and Functional Restoration of Sensory and Motor Systems”, University of Göttingen, Göttingen, Germany; Cluster of Excellence “Multiscale Bioimaging: from Molecular Machines to Networks of Excitable Cells” (MBExC), University of Göttingen, Göttingen, Germany; German Primate Center, Laboratory Animal Science Unit, Göttingen, Germany; German Center for Cardiovascular Research, Partner Site Göttingen, Göttingen, Germany; The University of Sydney, School of Medical Sciences (Neuroscience), Sydney, NSW, Australia; Else Kröner Fresenius Center for Optogenetic Therapies, University Medical Center Göttingen, Göttingen, Germany

## Abstract

A standard circuit motif in sensory systems is the pooling of sensory information from an upstream neuronal layer. A downstream neuron thereby collects signals across different locations in stimulus space, which together compose the neuron’s receptive field. In addition, nonlinear transformations in the signal transfer between the layers give rise to functional subunits inside the receptive field. For ganglion cells in the vertebrate retina, for example, receptive field subunits are thought to correspond to presynaptic bipolar cells. Identifying the number and locations of subunits from the stimulus–response relationship of a recorded ganglion cell has been an ongoing challenge in order to characterize the retina’s functional circuitry and to build computational models that capture nonlinear signal pooling. Here we present a novel version of spike-triggered non-negative matrix factorization (STNMF), which can extract localized subunits in ganglion cell receptive fields from recorded spiking responses under spatiotemporal white-noise stimulation. The method provides a more than 100-fold speed increase compared to a previous implementation, which can be harnessed for systematic screening of hyperparameters, such as sparsity regularization. We demonstrate the power and flexibility of this approach by analyzing populations of ganglion cells from salamander and primate (marmoset) retina. For the marmoset retina, we find that the sets of subunits obtained from analyzing populations of midget or parasol ganglion cells, respectively, form separate mosaics that tile visual space. Moreover, these subunit mosaics show alignment with each other for ON and OFF midget as well as for ON and OFF parasol cells, indicating a spatial coordination of ON and OFF signals at the bipolar-cell level. Thus, STNMF can help reveal organizational principles of signal transmission between successive neural layers.

## INTRODUCTION

In sensory pathways, neuronal signals typically carry spatiotemporal information about the outside world. The information is transmitted by an array of neurons, each encoding a constrained region of the stimulus space, defining its receptive field. A postsynaptic neuron can pool signals across multiple neurons in this array, constituting a receptive field that is larger than in the presynaptic layer. Owing to synaptic rectification or other signal transformations in the signal transmission, the integration of presynaptic signals is often nonlinear, which gives rise to substructure within the receptive field, for example, in the form of functional subunits that together compose the receptive field. Nonlinear integration of subunit signals then sets the stage for complex computations, as illustrated by examples from the retina (Zapp et al. 2022), primary visual cortex (Almasi et al. 2020; Bartsch et al. 2022; Lehky et al. 1992; Vintch et al. 2015), medial superior temporal visual cortex (Beyeler et al. 2016; Mineault et al. 2012), and the auditory system (Ahrens et al. 2008; Harper et al. 2016; Keshishian et al. 2020; McFarland et al. 2013).

In the retina, the neural network where vertebrate visual processing starts, subunits of ganglion cell receptive fields convey sensitivity of the cells to spatial contrast or to different types of motion stimuli (Demb and Singer 2015; Gollisch and Meister 2010; Kerschensteiner 2022). These ganglion cell subunits are thought to correspond to presynaptic bipolar cells (Borghuis et al. 2013; Demb et al. 2001; Liu et al. 2017). Their excitatory signals may be integrated nonlinearly by the ganglion cells (Borghuis et al. 2013; Demb et al. 2001), which plays a substantial role in sensitivity to high spatial frequencies (Enroth-Cugell and Robson 1966; Schwartz et al. 2012; Victor and Shapley 1979) as well as in local dynamics like contrast adaptation (Jarsky et al. 2011; Manookin and Demb 2006; Rieke 2001). Understanding how subunits compose the receptive field and how their signals are integrated can thus, on the one hand, provide essential information about the functional connectivity and the propagation of information in sensory networks, like how ganglion cells pool signals over an array of bipolar cells. These insights may then aid in explaining how ganglion cell receptive fields implement specific neural computations. On the other hand, identifying the layouts of subunits for different neuronal cell types can complement morphological findings by identifying differences in connectivity patterns and shared connections. In particular, the subunits obtained from the analyses of ganglion cells of a single type may be expected to jointly form a mosaic-like arrangement, covering retinal space by the set of subunits with little overlap between individual subunits. On the level of receptive fields, such mosaics are commonly observed for retinal cells of a given type, as has been reported across species for photoreceptors (Marc and Sperling 1977), bipolar cells (Cohen and Sterling 1990a, 1990b; Sterling et al. 1988), and ganglion cells (Anishchenko et al. 2010; DeVries and Baylor 1997; Gauthier et al. 2009).

The retina subjected to light stimulation has proved to be a suitable model for investigating the functional connectivity in sensory systems. Nevertheless, direct experimental investigations of the neuronal connections that constitute the subunits in the functioning network, for example by simultaneously stimulating and recording pre- and postsynaptic neuronal layers, remain challenging. And even though anatomy-based connectomics analyses have begun revealing the presynaptic networks of retinal ganglion cells (Bae et al. 2018; Helmstaedter 2013; Sigulinsky et al. 2024), limited sample sizes and the difficult task of detecting true connections have impeded obtaining a complete mapping of (excitatory) inputs, especially for larger ganglion cells, and connecting this to visual stimulus encoding. In light of the experimental limitations, several methods have been proposed to infer subunits computationally from recorded ganglion cell responses to visual stimuli. The methods include data-driven fitting of cascade networks that model subunits as linear filters with a nonlinear transfer function converging onto a ganglion cell (Freeman et al. 2015; Maheswaranathan et al. 2018; McFarland et al. 2013; Shi et al. 2019). Similarly, convolutional neural networks, which offer a comparable model of signal convergence with potentially deeper layering have been used to capture how local processing inside the receptive field of a ganglion cell shapes the cell’s visual responses (Maheswaranathan et al. 2023; McIntosh et al. 2016; Tanaka et al. 2019).

On the other side of the spectrum, there are statistical analyses of a cell’s stimulus-response relationship, such as spike-triggered clustering (Shah et al. 2020) and spike-triggered non-negative matrix factorization (STNMF; Liu et al. 2017). For example, STNMF relies on visual stimulation with spatial or spatiotemporal white-noise patterns and on analyzing correlations in the set of those stimulus patterns that elicited spikes (spike-triggered stimulus ensemble), while placing a non-negativity and a sparsity constraint on the extracted structures. Subunits identified with STNMF from salamander ganglion cells have been shown to match receptive fields of simultaneously recorded bipolar cells (Liu et al. 2017). However, the proposed implementation is computationally expensive, requiring on the order of several hours of analysis time per ganglion cell on a standard desktop computer. This has limited the control over the effects of hyperparameters, such as the strength of sparsity regularization, which needs to be adjusted to the right range for obtaining spatially localized subunits.

Here, we develop a new version of STNMF that addresses the shortcomings of previous implementations and investigate its applicability on recorded spike trains of ganglion cell populations from both salamander and primate retina. We introduce a novel combination of state-of-the-art algorithms of non-negative matrix factorization (NMF) and complement it with guided initialization procedures to improve speed and reliability. The resulting method allows subunit recovery for a cell in a matter of seconds on a standard desktop computer. We demonstrate how clustering consensus methods can be applied for hyperparameter selection and offer tools that reliably provide suitable values for sparsity regularization. This makes STNMF more versatile across cells of different functional classes and across datasets of different species. Finally, we use STNMF to investigate mosaic arrangements of subunits from the ON- and OFF-type pathways in primate retina, demonstrating that these are spatially aligned relative to each other, in contrast to the anti-alignment of ganglion cell receptive field mosaics (Roy et al. 2021).

## RESULTS

Receptive-field subunits are spatial regions that act as functional, nonlinear building blocks of a neuron’s receptive field. The basic conceptual model here is that subunits act as parallel linear stimulus filters whose outputs are nonlinearly transformed by the subunit nonlinearity and then combined by a weighted summation, potentially followed by another nonlinear transformation, the output nonlinearity (Zapp et al. 2022). STNMF aims at identifying these subunits by evaluating the statistical structure of spike-eliciting stimulus segments, the spike-triggered stimuli, under white-noise stimulation and is thus a type of spike-triggered analysis. Spike-triggered analyses have long been used for assessing receptive fields via computation of the spike-triggered average (STA; Bryant and Segundo 1976; Chichilnisky 2001; de Boer and Kuyper 1968) as well as for obtaining multiple (typically temporal) stimulus filters via spike-triggered covariance (STC) analysis (Cantrell et al. 2010; Fairhall et al. 2006; Gollisch and Meister 2008; Samengo and Gollisch 2013; Schwartz et al. 2006). As an extension of these approaches, STNMF uses statistical patterns (e.g., correlations) in the set of spike-triggered stimuli to obtain a parts-based (Lee and Seung 1999) decomposition of the receptive field.

To achieve this decomposition, STNMF seeks to approximate each of the spike-triggered stimuli by a linear combination of a limited number of basis patterns, or modules. Importantly, these modules are optimized to achieve the best possible approximation under the constraint that all their elements are non-negative. The goal of this analysis is to capture the shape and location of the subunits by individual modules, which is based on the following intuition. The limited number of modules, much smaller than the number of elements, say pixels, in the spike-triggered stimuli, forces the modules to incorporate the statistical structure of the spike-triggered stimuli, e.g., by letting pixels with strongly correlated activation across the spike-triggered stimuli be jointly represented within the same module. Furthermore, pixels within the same subunit generally have a higher correlation across the spike-triggered stimuli than pixels from different subunits because subunit nonlinearities typically induce a threshold effect. The threshold lets “negative” activation of one pixel counteract “positive” activation of another pixel more effectively if the pixels are part of the same subunit, making such patterns with opposing activation in the same subunit less effective in generating spikes and thus less likely to occur among the spike-triggered stimuli. Finally, the non-negativity constraint enforces a strictly additive composition of spike-triggered stimuli, rather than a combination where positive and negative parts of different modules can cancel out, which leads to sparse, localized modules, as desirable for subunit identification.

Mathematically, the approximation of the spike-triggered stimuli by a linear combination of the modules corresponds to a matrix factorization, in which the data matrix of all spike-triggered stimuli is approximated by a product of two smaller matrices, one containing the modules, the other the weights used for the linear combination. Finding an optimal factorization that best approximates the data matrix under the non-negativity constraint is known as non-negative matrix factorization and generally approached by iteratively improving the two matrices, as explained in more detail below.

One useful simplification for identifying spatial subunits of retinal ganglion cells is to reduce spatiotemporal spike-triggered stimuli to purely spatial stimuli by integrating out time. Since subunits of a given cell typically share the same temporal dynamics (Maheswaranathan et al. 2018), this has been a viable option to reduce input dimensions (Liu et al. 2017; Shah et al. 2020). To do so, the cell’s temporal filter, obtained as the temporal component of the STA, is used to compute a weighted spatial component for each spike-triggered stimulus, that is, a single effective spike-triggered spatial pattern (Kaardal et al. 2013; Liu et al. 2017).

Concretely, our analysis starts with recording spikes of retinal ganglion cells under spatiotemporal white-noise stimulation, organized in a checkerboard layout with light intensities of the checkerboard squares flickering randomly and independently. Similar to other spike-triggered analyses, the white-noise stimulus statistics are a prerequisite for the method so that statistical regularities in the spike-triggered stimuli are solely a result of the receptive field and its subunit structure. From the recorded data, we extract the spike-triggered stimuli for each cell, that is, the stimulus sequences preceding a spike over a predefined time window of 700 ms, covering the temporal sensitivity of the neuron. We compute a cell’s STA as the average of all its spike-triggered stimuli and separate it into a spatial and a temporal component (see Methods). For each spike-triggered stimulus sequence, we then take the average of the stimulus frames weighted with the temporal component. Furthermore, to focus the analysis on the relevant region in space, the time-averaged spike-triggered stimuli are cropped to roughly the region of a cell’s receptive field, here done by selecting the smallest rectangle containing the three-standard-deviation ellipse of a Gaussian function fitted to the spatial component of the STA.

The collection of these time-averaged, cropped spike-triggered stimulus patterns gives us the effective spike-triggered stimulus ensemble (STE) for a given cell. We arrange the STE into a pixel-by-spikes matrix (**Figure 1A**), the data matrix, which we also call the STE matrix, where each column collects all the pixel values of the effective spike-triggered stimulus for a given spike. We then perform semi-non-negative matrix factorization (semi-NMF; Ding et al. 2010) on this STE matrix. The use of semi-NMF, rather than a full NMF with non-negativity constraints on all matrices, enforces non-negativity only on one of the two matrices in the factorization. This accommodates the negative contrast values contained in our STE matrix.

**Figure 1.**
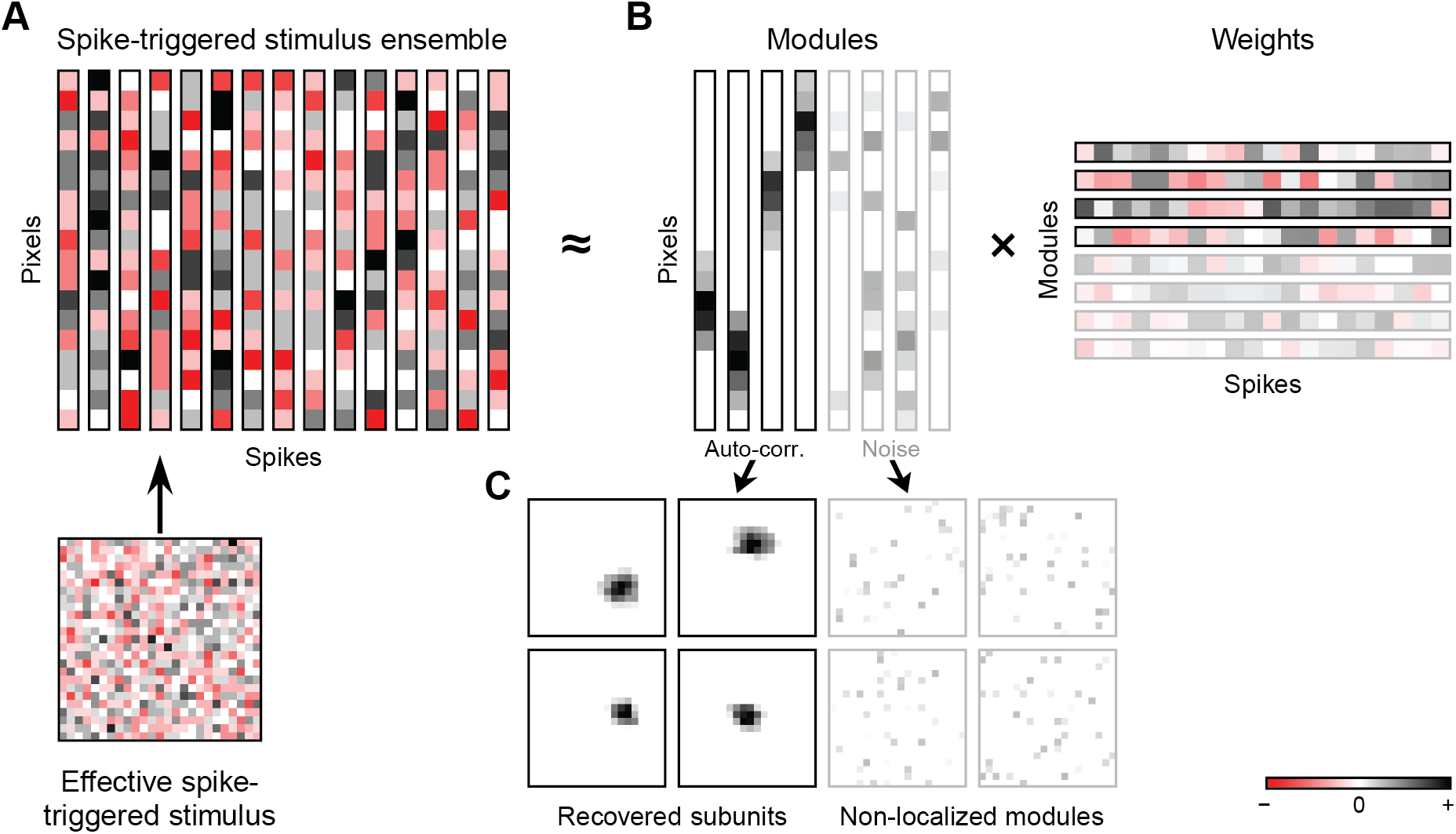
Conceptual sketch of spike-triggered non-negative matrix factorization. (**A**) The spike-triggered stimulus ensemble (STE, top) is a matrix of pixels by spikes. Each column is the effective spike-triggered stimulus of a given spike (bottom). The STE serves as the input to the matrix factorization (dimensions not to scale). (**B**) Using semi-NMF, the STE matrix is decomposed into two smaller matrices, the non-negative spatial modules and their corresponding weights. Together, these represent a lower-dimensional approximation of the STE. (**C**) The modules can be reshaped into two-dimensional spatial layouts. Some modules exhibit localized structure and are identified via their spatial autocorrelation as subunits, whereas others (here displayed with reduced saturation) capture noise.

The semi-NMF decomposes the STE matrix into two smaller matrices, the module matrix and the weight matrix (**Figure 1B**). The module matrix contains a given number of spatial modules, each corresponding here to one column of the matrix. The number of spatial modules is a free parameter in the analysis, which is chosen as explained below. The module matrix is optimized in the analysis to allow the best approximation of the STE matrix through a linear combination of the modules. The corresponding weights compose the weight matrix, with each column containing the modules’ weights that are needed for a given spike to best approximate the effective spike-triggered stimulus of that spike by a weighted sum of the modules. The optimization is here performed under a non-negativity constraint on the module matrix so that the modules emerge as purely additive building blocks. In addition, we apply a sparsity constraint to the module matrix, which aids the extraction of sparse, spatially localized modules.

The final modules typically come in two distinct types; they either have localized, patch-like structure or non-localized, noise-like speckles (**Figure 1C**). We automate the differentiation between these two cases by computing a measure of spatial autocorrelation (Moran’s I, see Methods) and identifying localized modules as modules with a Moran’s I value above a threshold (here taken as 0.25). These localized modules are taken as the recovered subunits within the receptive field of the ganglion cell. The non-localized modules, on the other hand, are typically dominated by pixels outside the receptive field, which do not contribute to activating the cell. The noise-like subunits therefore capture spurious correlations within the STE, essentially reflecting noise in the STE from finite sampling. For the detection of subunits, these noise-like modules can thus be ignored.

Prior to running STNMF, one has to select how many modules to use for the matrix factorization. In NMF, the optimal number of modules is generally not known a priori, and its selection poses a common hurdle. For STNMF, one might think that the number of modules should be equal to the actual number of subunits that compose the receptive field. This is not the case, however. Instead, we find, as previously described (Liu et al. 2017), that it is advantageous if the number of modules is somewhat larger than the number of (expected) subunits. The reason for this lies in the two distinct types of modules found in the STNMF solutions (**Figure 1C**). As the subunits are represented by a subset of the modules, namely those with localized structure, the additional modules are used by STNMF to capture noise in non-localized patterns.

Allowing for more modules in the STNMF analysis mostly increases the number of these non-localized modules, whereas the number of localized subunit-like modules remains largely unaffected, making the identification of subunits fairly robust to varying the number of modules. **Figure 2** provides example results for two salamander retinal ganglion cells, analyzed with STNMF using 15, 20, or 30 modules. This illustrates the either localized or noise-like structure of derived modules and that increasing module number mostly increases the number of the noise-like subunits, leaving the identified subunits approximately fixed. As previously suggested (Liu et al. 2017), we find 20 modules to be a good number for our application to retinal ganglion cells, at least in the datasets analyzed here, as the number of recovered subunits rarely exceeded ten. For other datasets, however, the number of modules might need to be adjusted, e.g., by trial and error to identify a range where localized subunits are robustly detected without depending subtly on the number of provided modules.

**Figure 2.**
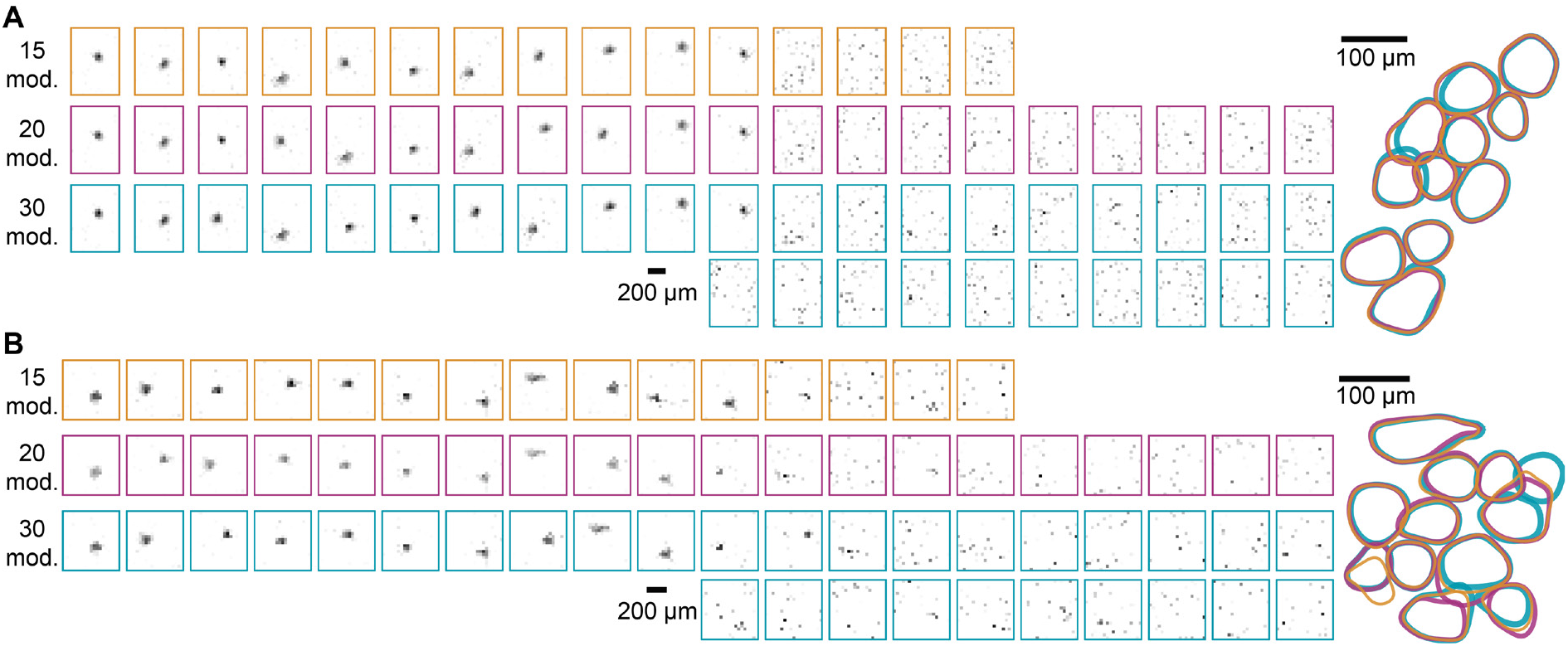
Effect of varying number of modules. (**A**) Modules extracted for a salamander retinal ganglion cell from an available dataset (Gollisch and Liu 2018), using either 15, 20, or 30 modules (mod.) in the STNMF algorithm. The outlines of localized modules, that is, the subunits, are shown overlaid for all the cases on the right. In this example, all three versions of STNMF recover the same subunits, and additional modules only contain noise-like patterns. (**B**) Same as (**A**) for a second sample salamander retinal ganglion cell. In this example, some deviations between the STNMF runs are apparent, e.g., an additional subunit found at the upper right by the 30-module version and a slightly shifted subunit in the lower left for the 15-module version. Such deviations occur in particular for subunits at the edge of the receptive field, reflecting reduced reliability for such subunits with typically smaller weight.

### Accelerated fast hierarchical alternating least squares speeds up subunit recovery

We investigated different NMF algorithms to provide the most suitable implementation in the context of subunit recovery. Finding exact NMF solutions is known to be a computationally difficult challenge because of its non-negativity constraint; in fact, it has been shown to be an NP-hard problem (Vavasis 2010), meaning that no algorithm exists for finding exact solutions where the computational time scales in a polynomial fashion with the amount of data. Thus, to obtain good approximate decompositions, NMF is generally performed by iterative improvements. Various algorithms facilitate these iterations under the non-negativity constraint. Many of them are accurate but computationally costly algorithms (Kim and Park 2011), as they often seek exact solutions at each iteration when updating either the modules or the weights and because they consider all data at once.

As an alternative, hierarchical alternating least squares (HALS; Cichocki et al. 2007; Ho 2008; Li and Zhang 2009) has gained popularity in recent years. HALS replaces accurate with approximated solutions at each iteration of NMF, but exploits the thereby gained iteration speed. Over more but faster iterations, it outperforms previous methods in many applications of NMF (Cichocki et al. 2007; Cichocki and Phan 2009; Gillis and Glineur 2012) while ensuring convergence (Gillis and Glineur 2008). Instead of inferring all modules from their weights at once, HALS updates the modules sequentially. This breaks down large matrix multiplications into smaller, more efficient vector operations. The speed of this approach compensates for this approximation because more iterations are possible in shorter time. Various modifications of the HALS algorithm have been proposed recently to further increase its speed or its accuracy.

Here, we combine two HALS algorithms into what we call Accelerated Fast HALS (AF-HALS). Fast HALS (Cichocki and Phan 2009) achieves a speed-up through computational simplification by employing vector normalization to cancel out terms in the update equation. This makes each NMF iteration less complex and thus reduces computation time. Accelerated HALS (Gillis and Glineur 2012) improves the factorization in a given iteration step by performing multiple consecutive HALS updates, as long as the number of floating point operations stays below that of the heavy matrix operations preceding the next iteration. This makes each iteration more accurate and thus reduces the required number of iterations. The algorithmic approach for the AF-HALS-based implementation of STNMF is depicted in a flowchart in **Figure 3**.

**Figure 3.**
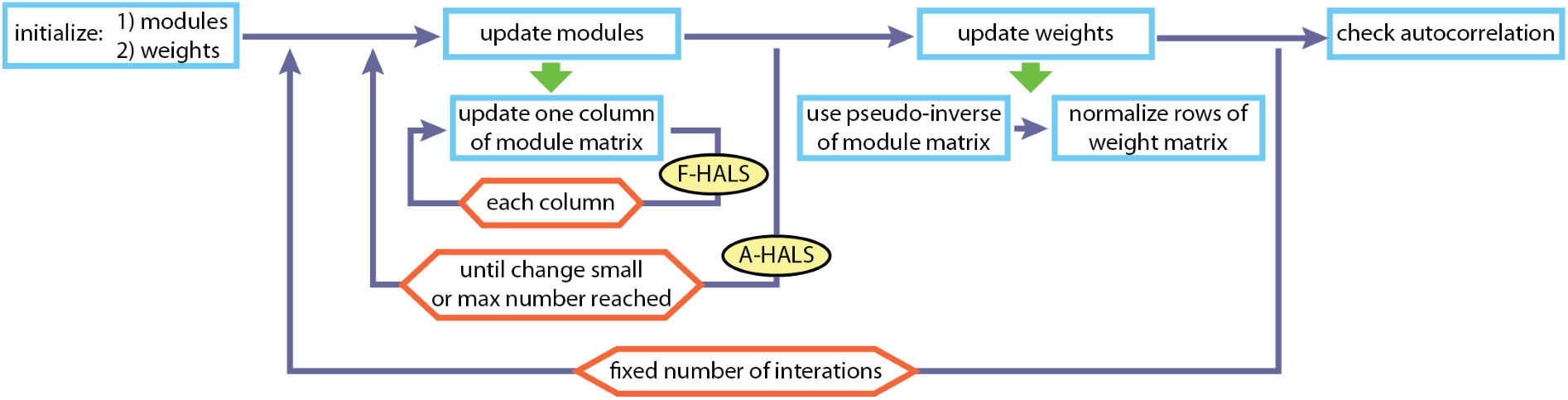
Flow-chart for the AF-HALS algorithm for STNMF. After initialization of the modules (either using random entries or by a guided initialization as explained below) and a subsequent computation of the initial weights, the main loop iteratively updates the module matrix and the weight matrix, here for a fixed number of iterations. The update of the modules consists of sequential column-wise updates (F-HALS loop), which are repeated several times (A-HALS loop). This allows re-using pre-computed matrix products several times while improving the module matrix estimate. The A-HALS loop stops when the change in the module matrix becomes small or when a maximum number of floating-point operations is reached. The update of the weights is an unconstrained optimization and therefore simply multiplies the pseudo-inverse of the current module matrix with the STE matrix. This is followed by a normalization of the rows of the weight matrix to avoid redundancies in the solution space. After concluding the iterations of alternating updates of modules and weights, subunits are determined as those modules (that is, columns of the module matrix) whose spatial autocorrelation (Moran’s I) surpasses a threshold value.

To demonstrate a direct comparison in speed and accuracy in the context of STNMF, we selected the popular active-set method (Lawson and Hanson 1974) as an alternative to AF-HALS. It is representative for current state-of-the-art NMF algorithms (Kim and Park 2011; Zhang et al. 2014) as it prevails for its relatively high accuracy at each iteration. The active-set method is based on optimal solutions to the sub-problem of updating one factorization matrix while keeping the other matrix fixed, though this optimal solution is approached iteratively in each step, thus also requiring a compromise between speed and numerical accuracy of the solution. A previous implementation of STNMF was based on the active-set method (Liu et al. 2017). We demonstrate the differences exemplarily on a retinal ganglion cell from the axolotl salamander retina, which had been recorded under white-noise stimulation as part of a previous study of subunit recovery with STNMF (Gollisch and Liu 2018; Liu et al. 2017).

We find that, using the same data, number of modules, and initial conditions, AF-HALS converges on localized modules much faster (**Figure 4**). Most importantly, the individual NMF iterations are considerably quicker with AF-HALS, reaching 10 iterations within a fraction of a second rather than the several seconds required by the active-set implementation, without sacrificing accuracy. For the first iterations, both methods, in fact, perform nearly identical steps, owing to the similarities in the objective of the iterations and to the identical initialization used here, which points both algorithms in the same direction for the initial improvement. As the optimization of the modules continues, the quality of the identified subunit structure remains similar or is even slightly better for AF-HALS, indicating that the approximation at each iteration is comparable to the numerical imprecision of the active-set method as implemented here. Thus, a clear subunit decomposition already emerges within the first few hundred iterations of AF-HALS-based STNMF, taking only a few seconds, with a remaining error that is smaller than what is reached by the active-set implementation even after several tens of minutes of runtime (**Figure 4**). In the end, both methods arrive at similar subunit estimates, but convergence for AF-HALS is reached much sooner by orders of magnitude in absolute computation time.

**Figure 4.**
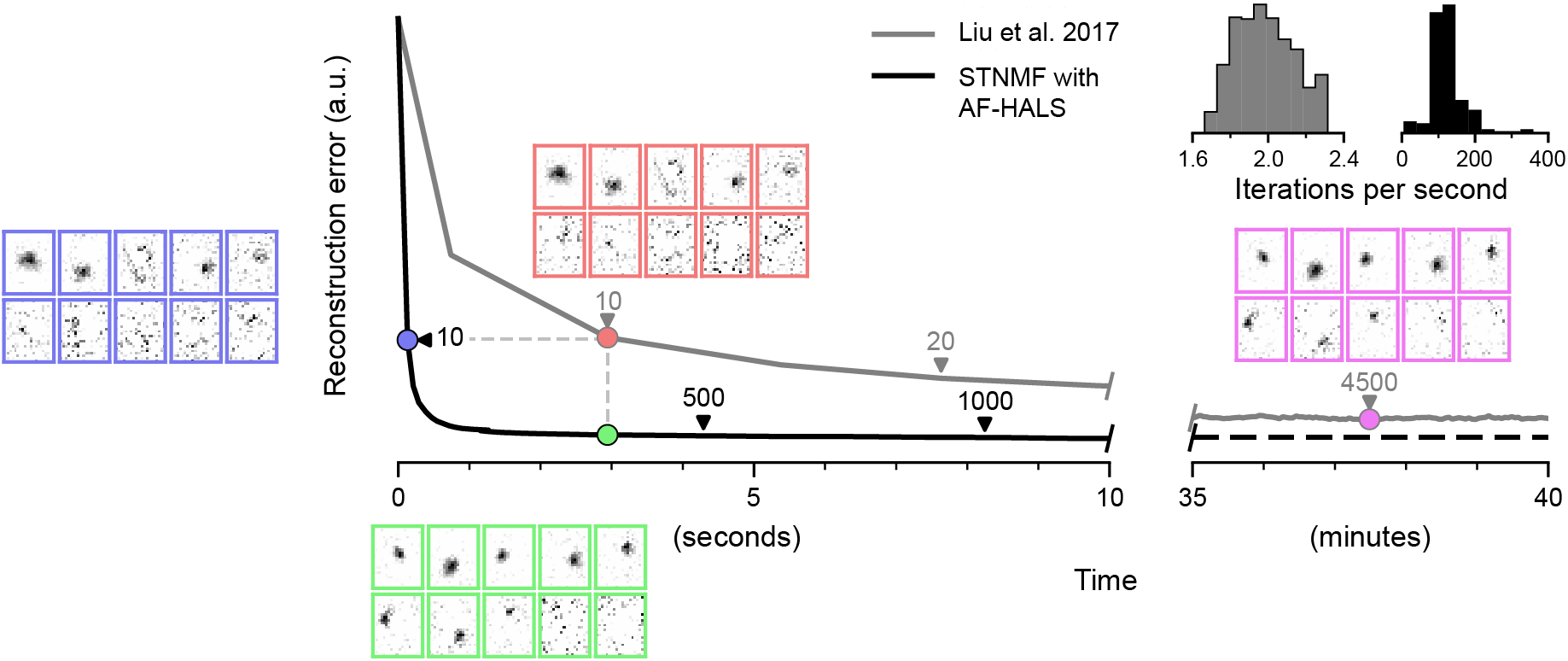
STNMF with AF-HALS converges faster than the previous implementation of STNMF. Reconstruction error of STNMF implemented with AF-HALS (black) and with the active-set method used by Liu et al. (2017; gray) for a sample salamander ganglion cell. Arrows denote the number of iterations. Insets allow visual inspection of the recovered modules at different states during the iterations. The reconstruction process is similar after ten iterations in both methods (blue and red insets), but the active-set method does not reach the low error that AF-HALS accomplishes within a few seconds. The visualized subunits of the active-set method after 45 minutes computation time (purple inset) arrive at the state that AF-HALS reached within five seconds (green inset). Top right: Distributions of iteration speeds (measured as inverse of each iteration duration) of the active-set method (gray) and of AF-HALS (black) for the sample cell.

### Guided initialization can replace repeated runs

As NMF is a non-convex optimization problem (Vavasis 2010), there are no guarantees for any given method to arrive at the globally optimal solution. In typical iterative implementations, solutions correspond to local optima and therefore depend on the starting point of the iterative search (Lee and Seung 1999), that is, the initial values of the modules. A popular way to compensate for the dependence on the starting point is to perform multiple repetitions of NMF starting from different random initializations of the modules and select the best of the obtained decompositions, the one with minimal residual reconstruction error (Berry et al. 2007; Cichocki et al. 2009).

However, random initialization generally does not offer a strong starting point for efficient convergence of the iterative procedures (Wild et al. 2004), and relying on many repetitions with different initializations poses a high demand on computational time. Furthermore, the selection of the best solution may depend on the particular measure of comparison. To circumvent these obstacles, we substitute the time-consuming repeated runs by implementing a guided initialization, based on singular value decomposition (SVD; Boutsidis and Gallopoulos 2008). Applying SVD to the matrix of the STE already yields structured components, akin to modules after several iterations of randomly initialized NMF, albeit with both positive and negative entries. Furthermore, SVD-based initialization has been shown to yield earlier convergence and smaller residual errors of NMF (Qiao 2015). To accommodate for the non-negativity constraint of NMF, methods of SVD-based initialization typically set negative values in the SVD components to zero before supplying these as initial conditions to the NMF algorithm.

As a further improvement, one can take advantage of the sign ambiguity of SVD and duplicate selected components with opposite sign before setting the negative values in all components to zero, as is done in non-negative singular value decomposition with low-rank correction (NNSVD-LRC; Atif et al. 2019). This offers potentially twice the amount of information from the selected SVD components. Furthermore, the procedure generates initial modules with around half of their values at zero, offering around 50% sparsity before even starting NMF. **Figure 5** illustrates this approach of module initialization for a sample salamander retinal ganglion cell. NNSVD-LRC subsequently uses the low-rank approximation of the input matrix constructed from the selected components to perform a few NMF iterations. The low-rank factorization reduces the computational complexity of the matrix multiplications involved, providing a computationally cheap head start for the full NMF. We modified NNSVD-LRC to be suitable for semi-NMF and improved its numerical stability (see Methods). This includes keeping the previously skipped duplicate of the component with the highest singular value and replacing the elements of columns whose entries are all zeros with a small non-zero constant to avoid convergence issues in HALS-based NMF.

**Figure 5.**
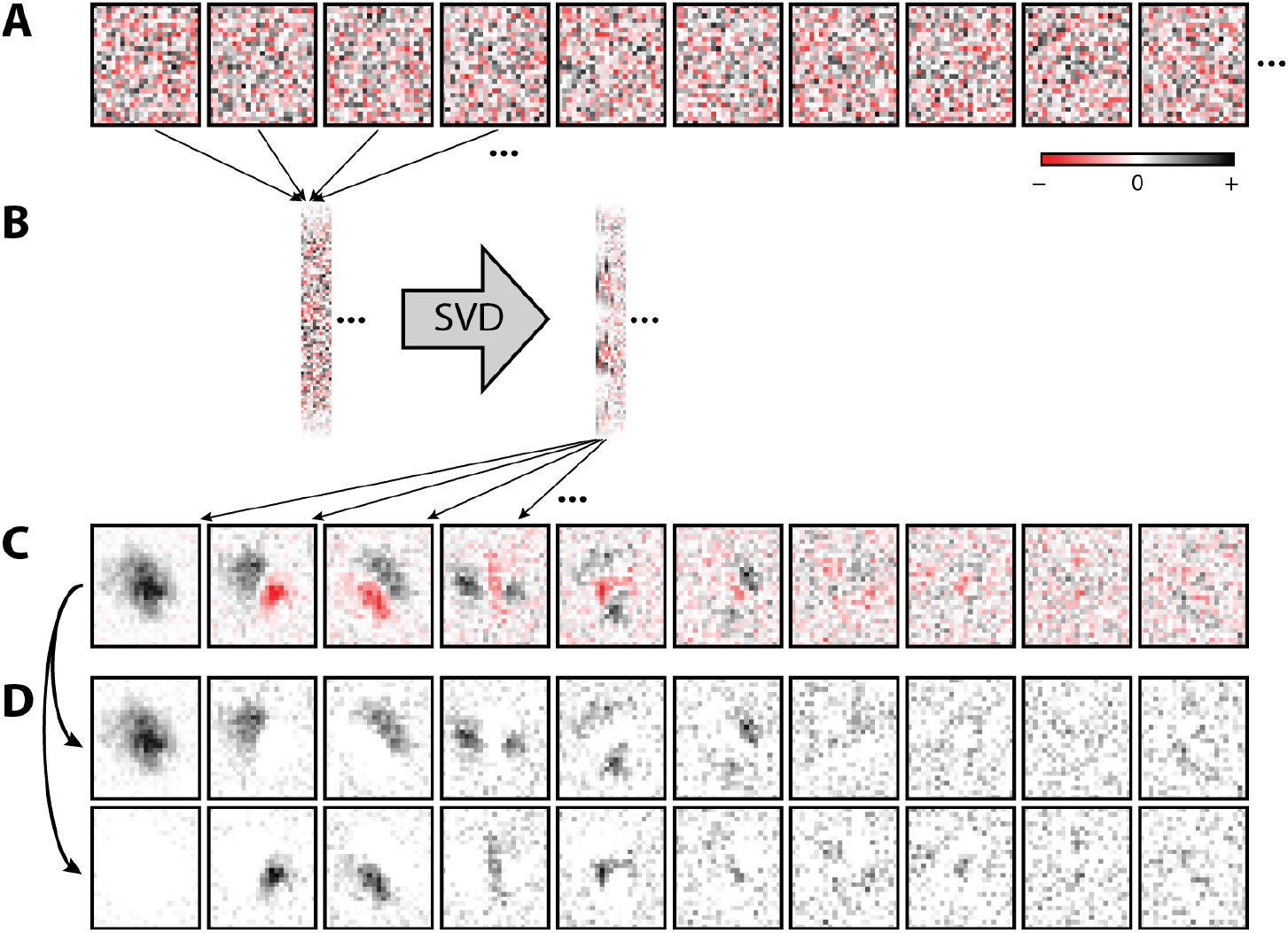
Example of module initialization with NNSVD-LRC. (**A**) Sample effective spike-triggered stimuli for a salamander retinal ganglion cell, here for the first 10 spikes in response to spatiotemporal white noise. (**B**) Arrangement of the effective spike-triggered stimuli into the STE matrix, which is then transformed by singular value decomposition (SVD). The transformed matrix is one of the matrices obtained from the SVD, scaled by the square-root values of the singular values. In the original STE matrix and the transformed version, every column contains the entries from a corresponding two-dimensional stimulus pattern, e.g., an effective spike-triggered stimulus for a column of the original STE matrix. (**C**) Display of the first ten spatial patterns contained in the transformed STE matrix, with each pattern corresponding to one column of the matrix after the SVD. (**D**) Construction of the 20 patterns used for initializing the modules. Each pattern from (**C**) is used twice, with a sign inversion for one of the copies, and negative entries are subsequently set to zero. To use these patterns as the initialization of the module matrix, the patterns are again flattened into individual columns of the module matrix. In all panels, black pixels correspond to positive numbers, red pixels to negative numbers, and white to zero.

We find that a single run of STNMF initialized with NNSVD-LRC can typically recover all recurring subunits that had been identified via 100 repetitions of randomly initialized STNMF. To show this, we compared STNMF results for an available dataset of salamander retinal ganglion cells (Gollisch and Liu 2018). We here focused on cells of the fast-OFF type, which are the most numerous and consistent in this dataset (Liu et al. 2017). For this comparison, we aimed at determining those spatially localized subunits that were robustly identified across multiple randomly initialized STNMF runs. We defined them as subunits that could be repeatedly identified across a majority of runs by their high spatial overlap (see Methods). We found that those robustly identified subunits matched the subunits recovered with NNSVD-LRC-based STNMF quite closely (**Figure 6**). This demonstrates the viability of the NNSVD-LRC initialization and supports the possibility of an additional 100-fold speed increase by omitting the repeated runs with random initialization, in addition to the already manifold speed increase obtained from applying AF-HALS.

**Figure 6.**
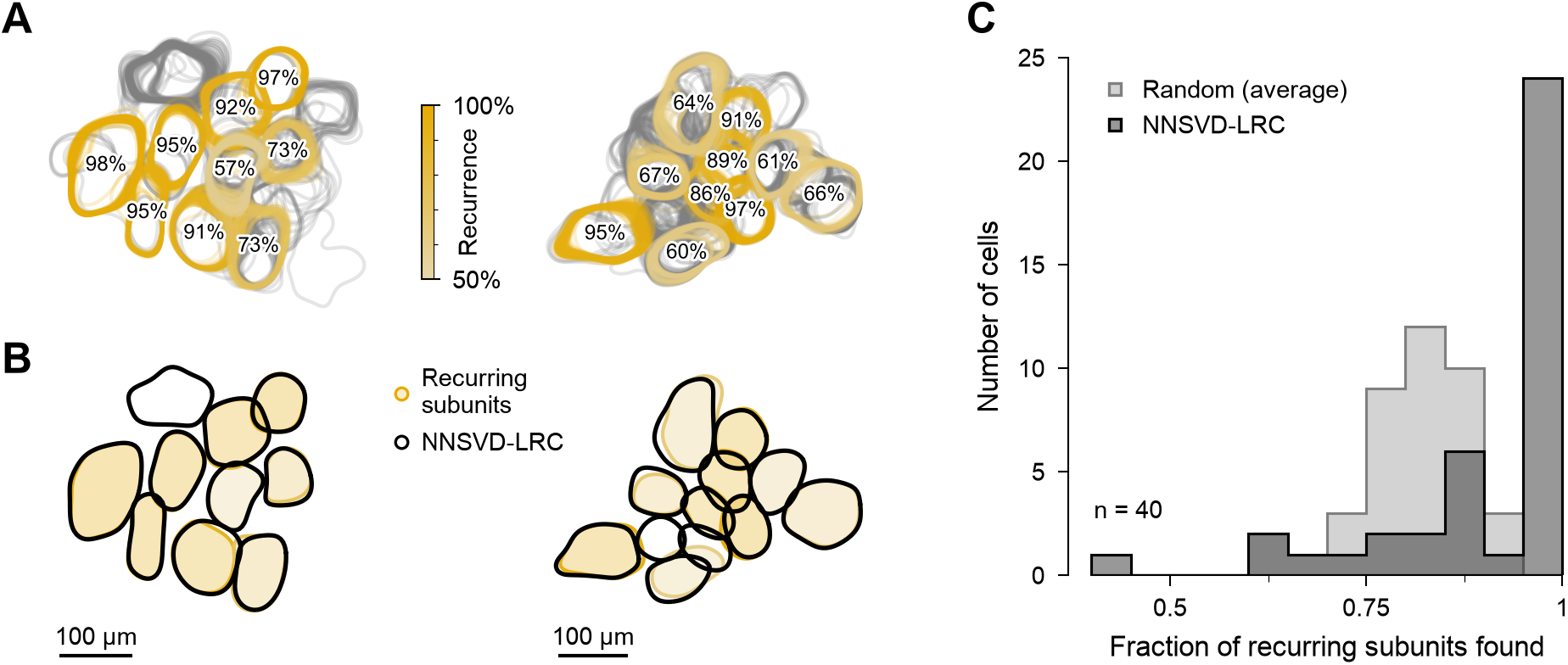
A single run of NNSVD-LRC-initialized STNMF yields subunits consistent with recurring subunits from 100 randomly initialized runs. (A) Contour outlines of subunits of 100 randomly initialized STNMF runs for two sample cells (fast-OFF type) from salamander retina. Subunits of different runs are regarded as identical if they exhibit substantial spatial overlap (Jaccard index > 0.5; see Methods). Subunits that emerged in more than half of the runs are colored, with saturation corresponding to the level of recurrence (see colorbar) and marked with a percentage of recurrence. (B) Comparison of subunit layouts recovered with NNSVD-LRC (gray) and with randomly initialized runs and a 50%-recurrence-criterion (yellow). Each yellow outline is the contour of the mean of the recurring subunits in (A). For the two sample cells, a single NNSVD-LRC-based STNMF run found all recurring subunits of the randomly initialized runs. (C) Distributions over 40 analyzed fast-OFF cells of the fraction of recurring subunits recovered by a single run with NNSVD-LRC (dark gray) or by 100 runs with random (light gray) initialization. Recovered subunits are determined via the Jaccard index (see above and Methods). NNSVD-LRC-based STNMF reliably finds more of the recurring subunits than the individual randomly initialized runs (averaged over 100 runs for each cell).

### Consensus analyses provide methods for tuning regularization in semi-NMF

In contrast to regular NMF, where the non-negativity constraint is placed on all matrices, semi-NMF only forces one matrix to be non-negative, in our case the modules, not the weights. Due to the loosened constraint, semi-NMF typically, without additional constraints beyond non-negativity of one factor matrix, fails to retain additive, parts-based solutions (Ding et al. 2010), that is, solutions where individual modules capture specific, often local, coherent components in the data rather than representing global, holistic features (Lee and Seung 1999). This crucial aspect of NMF can be recovered in semi-NMF through sparsity regularization on one of the factor matrices (Hoyer 2002, 2004; Kim and Park 2007, 2008). Unlike other methods of subunit recovery or machine learning in general, where regularization serves as an optional step to counteract overfitting or to alleviate the lack of sufficient data, sparsity regularization in STNMF based on semi-NMF is therefore an essential part of its functionality when parts-based decompositions are desired. To obtain localized subunits, we enforce the modules to be both non-negative and sparse. We achieve this by ℓ1-norm-based regularization, including a term in the objective function that penalizes large numbers of non-zero-pixel values in the modules.

As is customary, the strength of regularization is controlled via a regularization parameter. Increasing the value of the regularization parameter drives more pixel values in the modules to zero. Once regularization becomes too strong, subunits are forced to shrink in size or split into multiple smaller subunits until they eventually vanish. The optimal amount of sparsity depends on the dimensionality of the data. Accessible properties, like the number of pixels and number of spikes, as well as unknown characteristics prior to subunit recovery, like the true number of subunits and their sizes, play a role. Consequently, the regularization may have to be individually adjusted for cells that differ in their properties. On the other hand, for cells that share characteristics, similar strengths of regularization should be suitable.

The regularization requires the choice of a hyperparameter value to control the amount of sparsity. With our tractable implementation of STNMF, we here provide a method to do so in reasonable computation time. A common approach to finding the most suitable weight for regularization is through cross-validation. However, techniques that rely on held-out data are problematic in applications of unsupervised learning like NMF (Bro et al. 2008; Owen and Perry 2009). In STNMF, for example, held-out data would typically correspond to a subset of spikes (and thus columns of the STE matrix holding the corresponding effective spike-triggered stimuli). The module matrix could then be obtained without the held-out data, but evaluating how well the held-out part of the STE matrix can be reconstructed with the obtained modules still requires the corresponding part of the weight matrix. These weights are specific for the spikes in the held-out data and would therefore have to be recomputed for the test data. Despite the use of held-out data, this can cause overfitting-like effects and bias the evaluation on the test set by leading to artificially low reconstruction error of the factorization. Adjusted cross-validation-like procedures for matrix factorization have been proposed (Fu and Perry 2017; Owen and Perry 2009) based on selectively leaving out data entries across both rows and columns of the input matrix (pixels and spikes) and keeping these randomly scattered matrix elements as test data. However, using missing or masked entries in the input matrix does not generalize well to accelerated NMF algorithms like Accelerated HALS because the method’s efficiency relies on vectorized calculations and matrix multiplications that are factored out of the column-wise updates.

Instead of cross-validation, we therefore turn to the solution of a similar parameter-selection problem. For clustering techniques, a common hurdle is the choice of the optimal number of clusters. This problem also translates to NMF and may be addressed by using so-called consensus methods (Monti et al. 2003). To explore this approach to find a suitable regularization parameter for sparsity, we temporarily neglect the aforementioned SVD-based initialization and revert to repeated runs of NMF with different random initializations. As explained in more detail below, the variability in the solutions across runs is then exploited to find the most consistent decomposition among the sets of solutions. The more appropriate the choice of the sparsity regularization parameter, the more the extracted modules become similar across solutions (Brunet et al. 2004), which can be used as a criterion to optimize the regularization parameter.

### Consensus analysis allows selection of regularization parameters for STNMF

In the following, we demonstrate how consensus analysis can be used to optimize regularization for STNMF, based on the data recorded from ganglion cells of a salamander retina (Gollisch and Liu 2018). **Figure 7** illustrates the sequence of steps. As a start, it is often useful to first identify the rough range of regularization parameter values worth considering. We do this by running STNMF (with NNSVD-LRC initialization) and observing how the obtained sparsity, measured as the fraction of zero values in the modules, depends on the level of regularization (**Figure 7A**). This can be used to find an upper bound for the regularization parameter, beyond which nearly all values in the modules approach zero and which defines the range for subsequent, more detailed analyses. This initial range finding can be performed for individual cells or representative examples of a cell class and can be carried out with a shortened STNMF analysis. For the present data, we found that 200 iterations already suffice to estimate sparsity.

**Figure 7.**
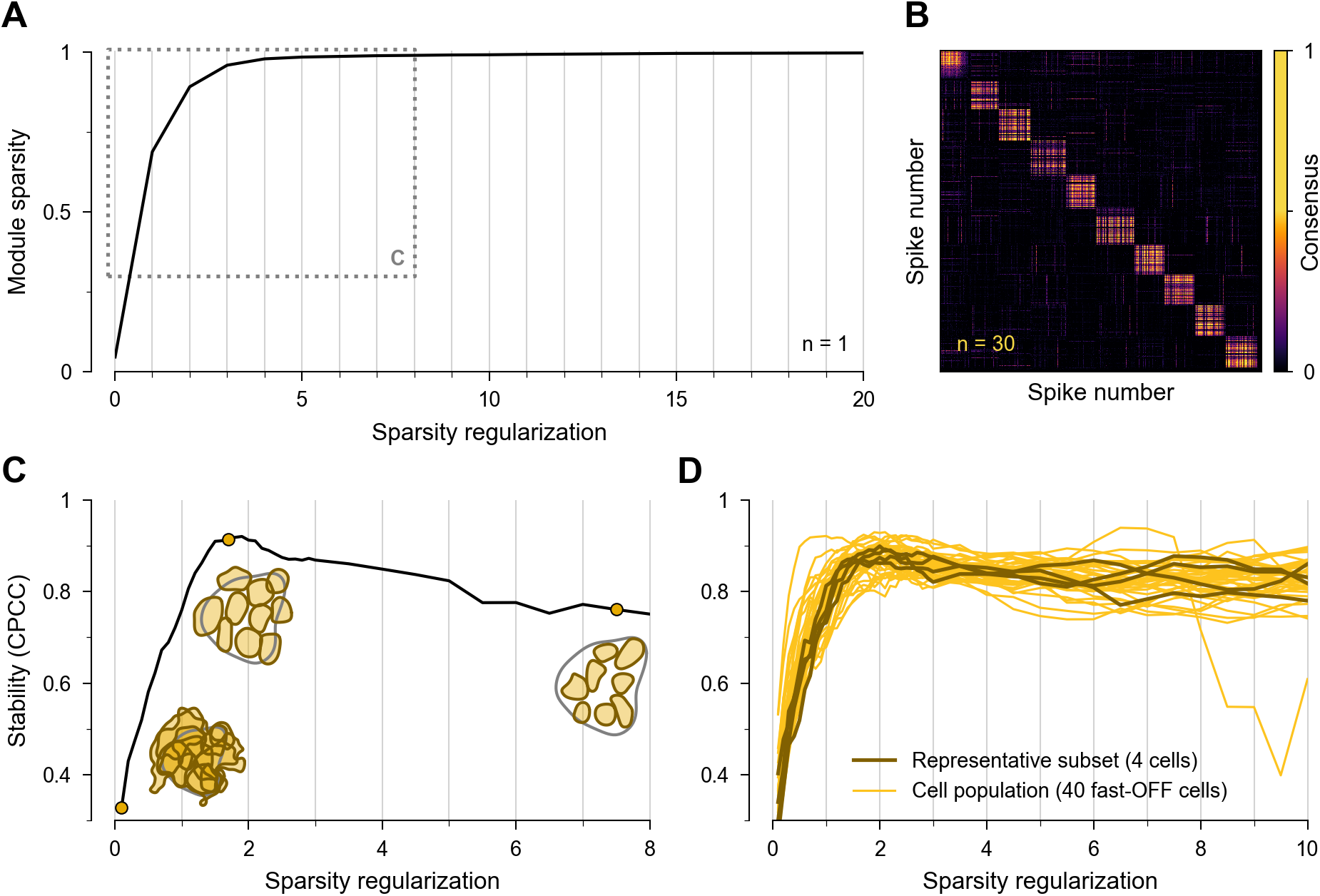
Consensus analysis for determining the weight of sparsity regularization. (**A**) Module sparsity (fraction of zeros in the modules) of the decomposition with increasing sparsity regularization for a sample salamander cell. The first step in the analysis is finding the window (dotted frame) in which regularization starts to dominate the decomposition, that is, module sparsity approaches unity. Within that range, the consensus of decompositions is probed. (**B**) Reordered consensus matrix showing the pairwise agreement between spike pairs across 30 decompositions. High consensus is indicated by the ten clusters on the diagonal corresponding to ten reliably recovered subunits as visualized in the second subunit layout inset of (**C**). (**C**) Stability curve for the same cell as obtained from consensus matrices at different regularization strength. The diagonal clustering structure in (**B**), corresponding to the stability of the assignment of spikes to subunits, is measured with a scalar value, the cophenetic correlation coefficient (CPCC). The CPCC increases with increasing sparsity regularization, along with refined subunit outlines (first versus second inset). After a peak in stability, subunits become too sparse (third inset) until they eventually vanish when regularization becomes too strong. (**D**) The stability curves for a wide range of regularization parameters of a subset of cells (dark) compared to the curves of all 40 fast-OFF cells (yellow) from a salamander retina.

For a given regularization strength in this identified range, we then assess the consensus of different solutions, obtained by running STNMF with different random initializations. In clustering, the consensus is typically analyzed via the consensus matrix, which contains for each pair of samples the fraction of times that the two samples were clustered together. In STNMF, the spikes take the role of the samples, and the weight matrix denotes their affiliation with the different modules, analogous to different clusters. This, however, is not a hard cluster assignment, but a soft weighting of how much each module contributes to reconstructing a given spike-triggered stimulus. Thus, to build the consensus across repeated runs, we determine the module most strongly affiliated with each spike as the module with the maximum weight for the spike. For our purpose, we further slightly modify the calculation of the consensus matrix by only allowing localized, subunit-like modules to contribute positively to the consensus. That is, if a pair of spikes is affiliated with the same non-localized module for a given STNMF run, this is treated like a case where affiliations do not match. We found that this adjustment reduced noise in the consensus analysis that likely comes from spurious coincidences of spikes with noise-like modules. The consensus matrix then contains, for all pairs of spikes, the fraction of randomly initialized STNMF runs for which the pair is affiliated with the same localized subunit (**Figure 7B**).

As proposed previously (Brunet et al. 2004), we quantify the dispersion of values in the consensus matrix by its cophenetic correlation coefficient (CPCC), which is a measure of the stability of the obtained subunits across random initializations. The cophenetic correlation measures the degree of clustering in the consensus matrix in a hierarchical clustering distance manner. This provides a scalar metric between zero and unity, with higher values indicating increased stability of the decomposition. Using the CPCC values, we then compare the stability of the subunit decomposition across different sparsity parameter values in the originally identified range (**Figure 7C**). We find that stability first increases with stronger sparsity regularization, as it progressively constrains the solution and reduces variability among the decompositions, and then plateaus or declines again when regularization becomes too strong.

It is also interesting to observe how the extracted subunit layouts vary in this range of different regularization strengths, as shown by the corresponding subunit layouts shown as insets in **Figure 7C**. Without sparsity regularization, we obtain modules that span the entire receptive field with strong overlap. As regularization strength increases, these modules first break up into large subunits that subsequently split into multiple smaller subunits. For a certain range, the subunit layout becomes more stable, as indicated by the maximal values of the stability curve, which coincides with a layout of similarly sized subunits that tile the receptive field. Around this region of peak stability, variations are mostly observed for subunits at the periphery of the layout. As sparsity regularization is further increased beyond this optimum of stability, some subunits are lost and others become smaller so that the receptive field is no longer covered by the subunit layout. Eventually, at the extreme end of strong regularization (not shown in **Figure 7C**), the reduction to very few or even only a single localized subunit can cause near perfect consensus (in particularly since our computation of the consensus matrix only considers localized subunits) and a corresponding rise in the stability curve. In practice, however, this extreme region can easily be avoided by observing the corresponding subunit structures.

The most important feature of the stability curve is that it exhibits a distinctive bend or peak after the initial sharp increase before it plateaus or decreases. The location of the bend or peak indicates a suited strength of sparsity regularization, providing a solution that is as stable as possible while affecting its structure as little as possible. Different techniques can be used to extract a specific regularization parameter value from the location, such as via a coefficient obtained from a fitted saturation curve. For our data, the bend or peak in the curves was readily identifiable and we simply selected a parameter value at this location by visual inspection. After determining a suitable regularization parameter, we run STNMF in that configuration initialized with NNSVD-LRC to obtain the final set of localized subunits.

### Suitable regularization parameters can be inferred from a subset of cells

We applied the consensus analysis to the salamander data of fast-OFF retinal ganglion cells. To obtain a suitable sparsity regularization parameter for subsequently identifying the subunits for all cells, we proceeded as follows. Among the population of fast-OFF cells, we selected a representative subset with respect to properties like receptive field diameter, number of spikes, and amount of noise in the STA. After initially determining a range of interest by observing the effect of regularization on module sparsity as in **Figure 7A**, we first investigated the solution stability for a coarse selection of regularization parameter values, performing only five repetitions at each sparsity value to save time.

This can be used to verify or further narrow down the range of regularization parameter values for the final analysis. In this range, we then performed a full consensus analysis at high resolution of regularization coefficients on the subset of cells (**Figure 7D**). We propose to apply between 20 to 30 repetitions for each regularization parameter value with 1,000 STNMF iterations or until convergence to obtain a reasonable estimate of stability. Finally, we extracted the suitable sparsity regularization parameter from the bend in the average of the stability curves. Beyond the subset of cells used for tuning the regularization, we then applied that regularization parameter to NNSVD-LRC-based STNMF to the entire population of cells to identify the final subunit layouts for all of them.

By applying consensus analysis to the full population over a wide range of sparsity regularization, we verified that parameters inferred from a subset of cells can be extrapolated to the entire population. For the sake of thoroughness, we analyzed a range of 44 sparsity parameter values between zero and ten, using a comparatively large number of repetitions (50) for each of them and an increased number of iterations (5,000) in the STNMF algorithm (**Figure 7D**, bright curves). Note, that this extensive analysis takes substantially more time than the procedure described in the paragraph above and was performed here for comparison purposes only. Occasionally, the stability curve can abruptly decrease and approach low values, as seen in **Figure 7D** for one example. This occurs due to excessive regularization strengths beyond the appropriate level, breaking subunits arbitrarily into smaller structures and thereby preventing consensus across repeated decompositions. Otherwise, the stability curves are generally well in agreement with each other so that an arbitrary subset of cells is representative of the population here.

### Subunits match previous analyses of bipolar cell receptive fields

We recovered subunits of the fast-OFF ganglion cells in the salamander data given the procedure described above. To estimate the subunits of all cells, STNMF took a total of five minutes on a conventional office computer averaging at around 9 seconds per ganglion cell. This is well beyond a 100-fold reduction in computation time compared to the earlier implementation based on the active-set method. Yet, the obtained subunits match the results from that previous analysis (Liu et al. 2017). We observe a close resemblance in the set of subunits of individual cells (**Figure 8A**) and find the subunit outlines to align for the cell population (**Figure 8B**).

**Figure 8.**
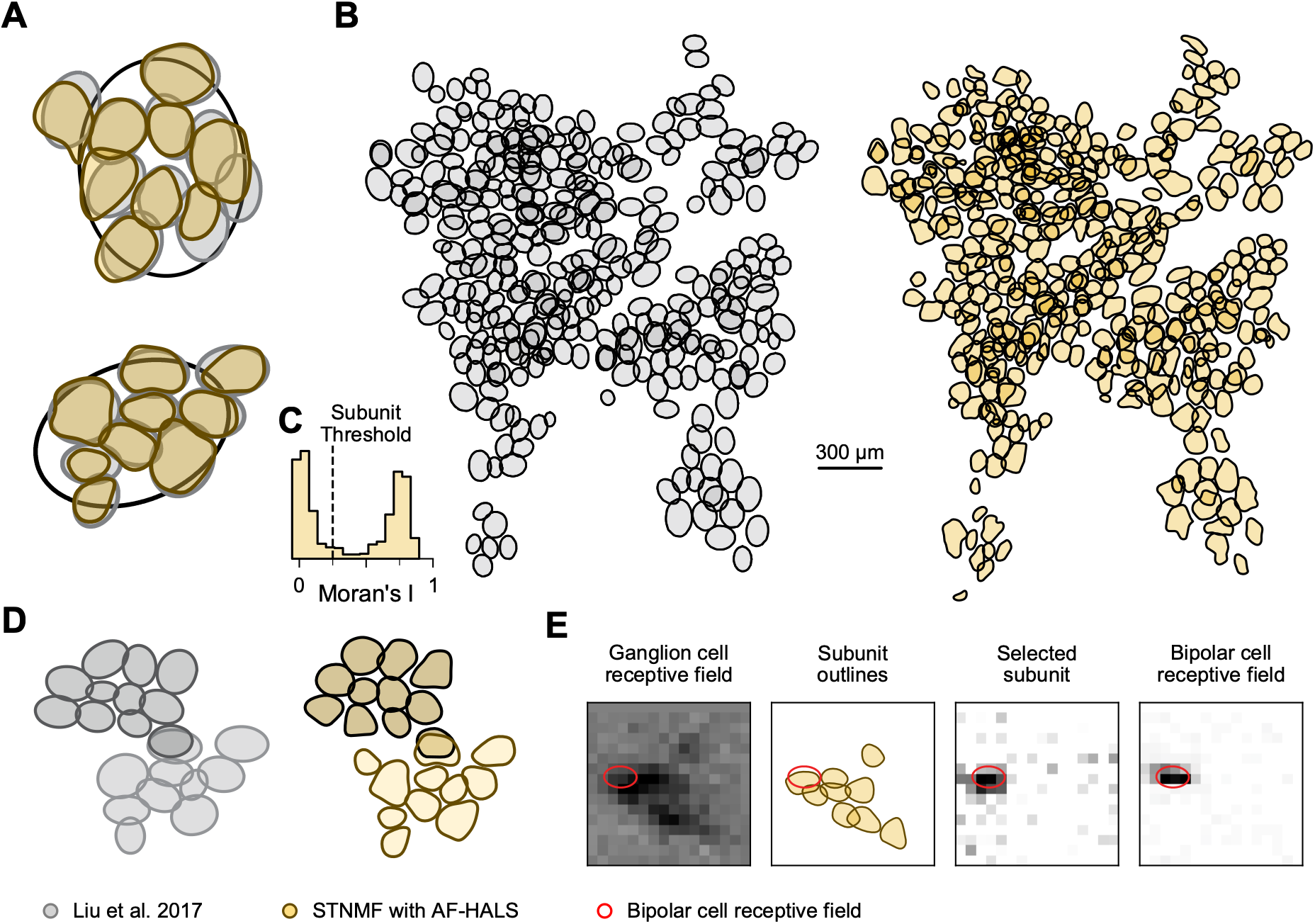
Subunits of AF-HALS-based STNMF match previous findings and bipolar cell receptive fields. AF-HALS-based subunit layouts of salamander cells (yellow) are compared to results from Liu et al. (2017; gray). Subunits are represented as 1.5-sigma ellipses of Gaussian fits in gray and as yellow contour lines (see Methods). (**A**) Decomposition for two example cell receptive fields. Receptive field outlines are 1.5-sigma ellipses of Gaussian fits. The AF-HALS-based subunit layouts resemble the previously estimated subunits. (**B**) Comparison of subunit mosaics of a population of fast-OFF cells, colors like in (**A**). Most of the previously identified subunits as well as some additional ones are recovered. (**C**) Distribution of the values of spatial autocorrelation, Moran’s I, for all modules obtained here with AF-HALS-based STNMF from the salamander dataset, displaying a strongly bimodal distribution. For consistency with the earlier analysis of the same data (Liu et al. 2017), subunits were defined as modules with Moran’s I larger than 0.25, though a slightly higher threshold might have better reflected the bimodal distribution. (**D**) Examples of subunit layouts for adjacent fast-OFF ganglion cells with overlapping (shared) subunits, as obtained with both methods. (**E**) Comparison of recovered subunits with the receptive field of a simultaneously recorded bipolar cell. One of the subunits (yellow outlines) matches the bipolar cell receptive field (red outline). Panels (**A** gray), (**B** left), (**D** left), and (**E** rightmost) are adapted from Liu et al. (2017) licensed under CC BY 4.0.

We also used this analysis of a large population of cells to check our criterion of distinguishing between localized modules corresponding to subunits and non-localized modules that capture noise in the STE. As explained above, we distinguished subunits from non-localized subunits by assessing the spatial autocorrelation, taking modules with a Moran’s I value of at least 0.25 as a subunit. Evaluating the distribution of values for all modules in this cell population confirmed the existence of two distinct types of modules with different spatial structure, as the distribution of Moran’s I values is strongly bimodal (**Figure 8C**). By identifying the location of the minimum between the two modes of the distribution, this bimodality can be used to objectively define the separation between modules defined as subunits and those considered noise. For comparability with previous work (Liu et al. 2017), however, we here kept the threshold at 0.25, which is close to but slightly smaller than the minimum.

Furthermore, we were able to reproduce previously reported overlaps between subunits across cell pairs (**Figure 8D**). For some of the receptive fields, we recovered additional subunits to the otherwise matching subunit layout that had not been detected before. The differences we observe typically concern subunits with relatively small average weights in the STNMF decomposition. One possible explanation for these unmatched subunits may be the differing procedures to determine the final subunit layout. In the present work, the subunits correspond to the localized modules from STNMF without further curation. In the previous analysis (Liu et al. 2017), the final subunits were obtained from the average of reoccurring subunits over multiple repetitions. Lower-weight subunits that occurred less frequently across the repetitions might have been more likely to be excluded by the previous method.

Experiments combining multi-electrode recordings of ganglion cells with single-cell recordings of bipolar cells had been performed previously in salamander retinas to investigate the relationship between computationally inferred subunits and measured bipolar cell receptive fields (Liu et al. 2017). Here, we confirm the results using the data of the respective ganglion cells. We find similar overlaps between subunits and bipolar cell receptive fields (**Figure 8E**). These findings suggest that our implementation can reveal bipolar cell receptive fields as subunits.

Another way to test whether the identified subunits reflect actual structure in the biological substrate is to perform the analysis for independent parts of the data and check whether consistent results are obtained. Here, we subdivided the recorded spikes into a first and a second half (leaving out ten spikes in the middle as a buffer to avoid any overlap from spikes occurring in the same time bin) and compared the STNMF results for these independent data sets. As shown by two examples in **Figure 9**, many derived subunits are virtually identical for the two data halves and also match the subunits obtained by analyzing the set of all spikes.

**Figure 9.**
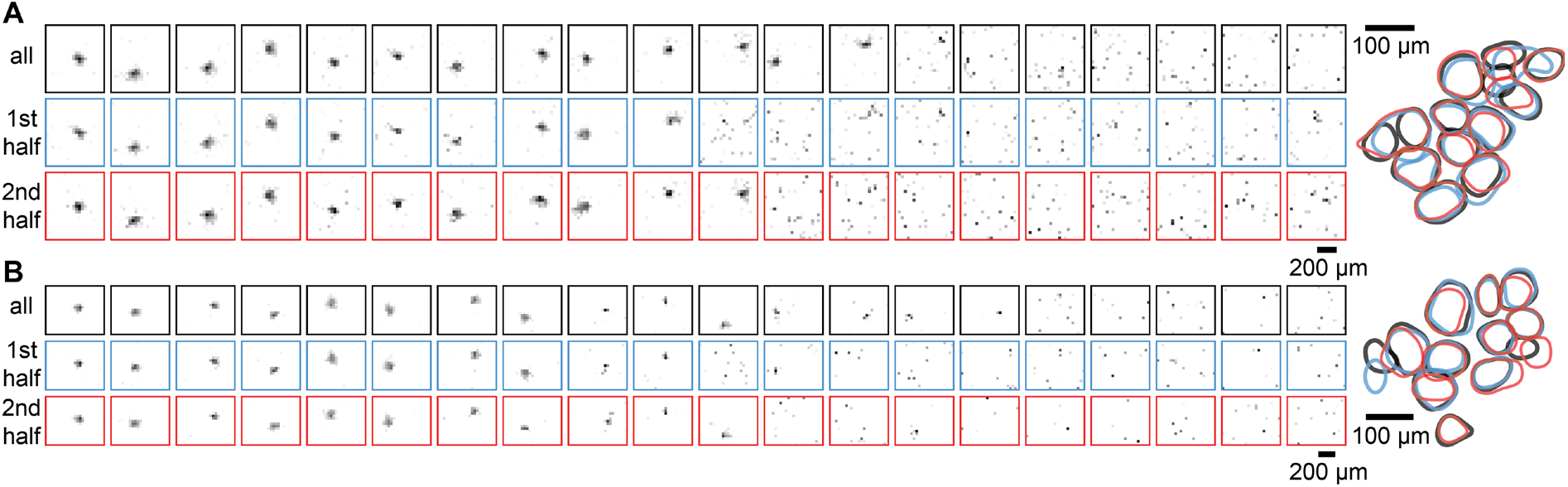
Comparison of identified subunits by STNMF from separate halves of the data. (**A**) Modules extracted for a salamander retinal ganglion cell, using either all available spikes (top, black rectangles), the first half of the spikes (middle, blue rectangles), or the second half of the spikes (bottom, red rectangles). The outlines of identified subunits (localized modules) are shown overlaid on the right. (**B**) Same as (**A**) for a second sample salamander retinal ganglion cell.

Some deviations occur primarily for subunits near the edges of the receptive fields, where subunit weights are presumably smaller and the subunits’ influence on spike generation thus weaker. Here, individual subunits might be missed by analysis of one of the halves, or the different analyses might disagree on whether certain subunits should be split or joined together. Overall, however, the close match of many subunits as obtained by independent parts of the data corroborate that the method extracts actual biological features within the receptive fields, and bipolar cells remain the simplest and most obvious interpretation.

### STNMF recovers subunit mosaics for all of the four major cell types in the primate retina

With the new, fast implementation of STNMF in place, we next aimed at performing subunit analyses of ganglion cell populations in the primate retina in order to demonstrate how STNMF can provide insight into the subunit organization of different ganglion cell types. To this end, we recorded spiking activity from ganglion cells in isolated marmoset retina under white-noise stimulation. We identified the four major cell types of the primate retina, ON and OFF parasol as well as midget cells, according to their receptive field sizes and temporal filters (see Methods). STNMF was able to identify subunits from the spiking data of individual cells for each of the four cell types (**Figure 10A**). Furthermore, the separation of the cells into four distinct types with at least partial tiling of receptive fields (**Figure 10B**) allowed us to identify the layout of subunits on a cell-type-specific population level (**Figure 10C**). By doing so, we found not only that subunits of individual ganglion cells cover the cell’s receptive field area with little overlap but also that subunits associated with different ganglion cells of the same functional type intertwine nearly seamlessly so that it is not easily visible where the boundaries between ganglion cell receptive fields are. Thus, the whole set of subunits obtained from ganglion cells of the same functional type tile the retina and thereby form a subunit mosaic, akin to the receptive field mosaic that covers the retina with the receptive fields of a ganglion cell type (DeVries and Baylor 1997; Field and Chichilnisky 2007).

**Figure 10.**
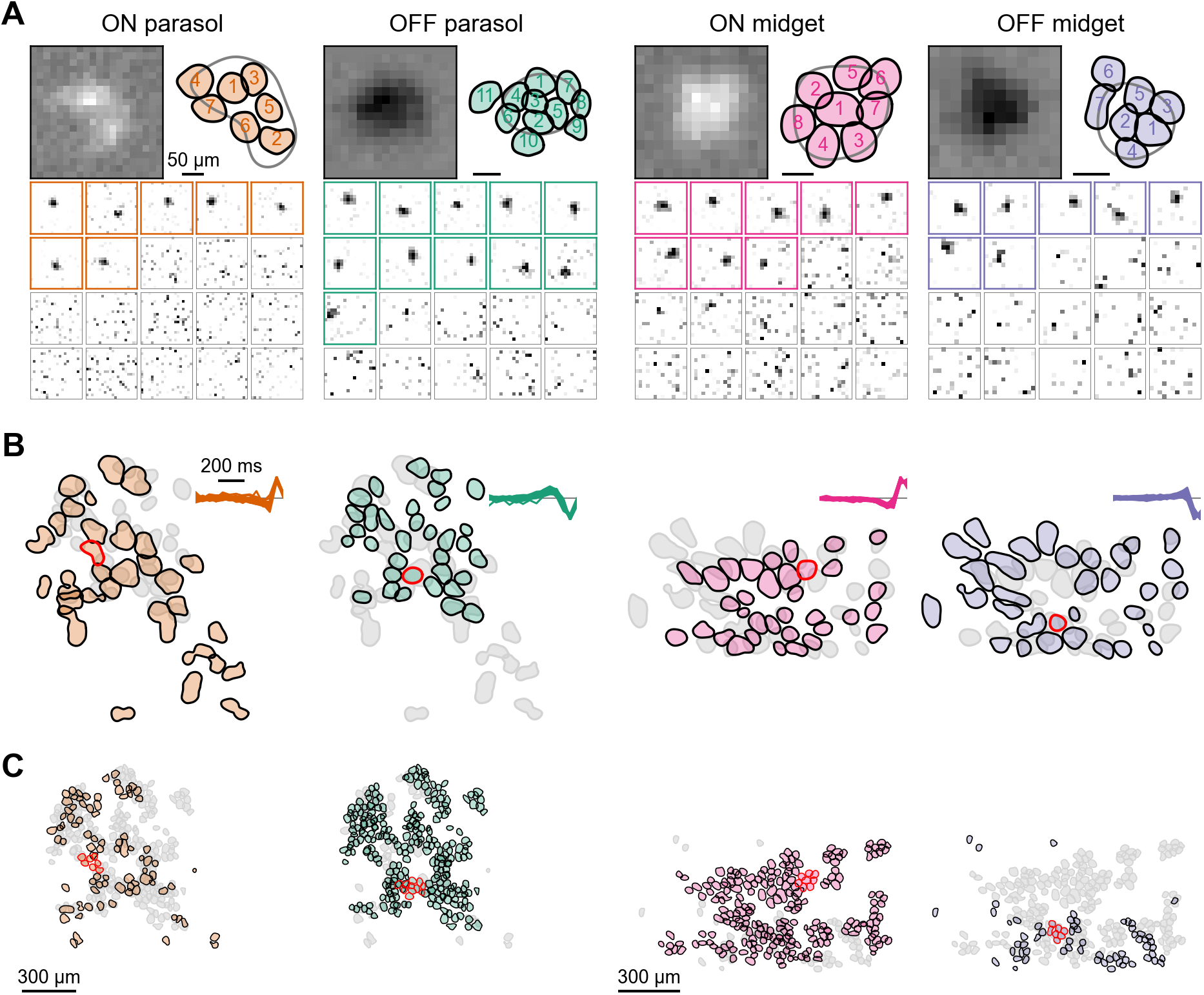
STNMF recovers subunits in all four major cell types of the marmoset retina. Data is shown from ON and OFF parasol cells (one retina; mid-periphery) and ON and OFF midget cells (one retina; periphery). Note that the scale bars differ between the four sample cells. (**A**) Receptive fields, STNMF modules, and subunit layouts of sample ON and OFF parasol cells and ON and OFF midget cells. Modules identified as localized subunits by their autocorrelation (see Methods) are marked by colored frames and depicted together in a colored subunit layout of contours superimposed on the receptive field contour (gray; top right). The outlines are numbered in descending order according to the mean STNMF weight. (**B**) Receptive fields of ganglion cell populations for the four main types (colored). Gray outlines correspond to the receptive fields of the corresponding cell type with opposite polarity of preferred contrast. The temporal dynamics of the receptive fields are depicted by the temporal filters of the spike-triggered average (insets; scale bar 200 ms). The sample cells of (**A**) are marked in red. (**C**) Subunit mosaics obtained from the ganglion cell populations in (**B**). Sample cells marked in red. For some cells (about one quarter to half of the OFF midget cells, 0–10% in other cell types), no subunits could be reliably identified. Scale bars (300 µm) correspond to both (**B**) and (**C**).

The availability of four distinct cell-type populations also allowed us to investigate the generality of the consensus analysis and to probe whether optimal sparsity regularization is cell-type-specific. As cells of the same functional type share properties like receptive field size and firing rate more so than cells of different types, the required sparsity may differ across cell types. To analyze whether this is reflected in the optimal sparsity regularization parameter, we examined the regularization-dependent consensus and compared the resulting stability curves for a wide range of regularization parameter values. For each of the four types, we found stability curves as expected, with a bend that indicates the optimal regularization strength (**Figure 11A**).

**Figure 11.**
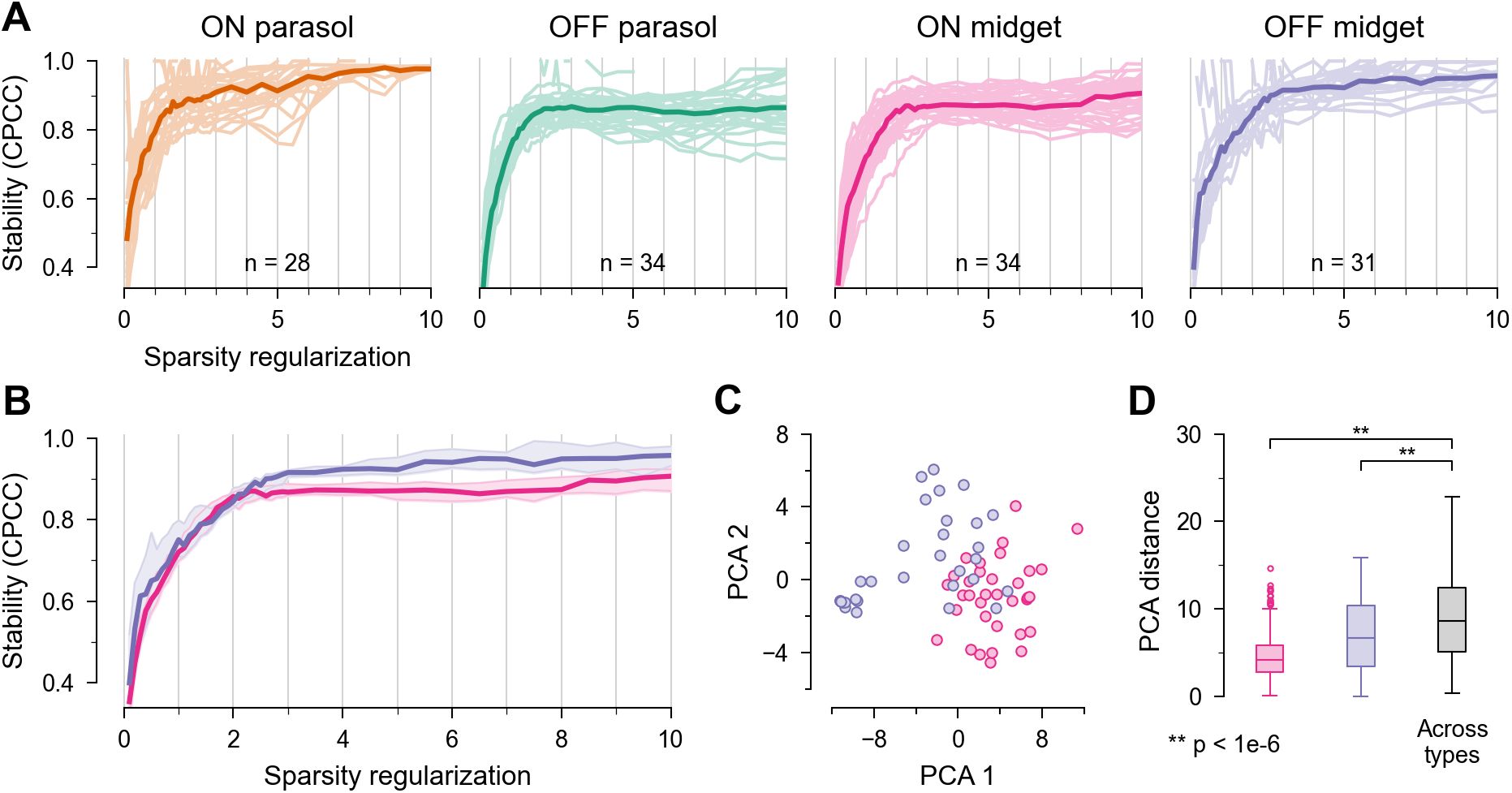
Stability curves align well among cells of same functional type. (**A**) Stability defined by the cophenetic correlation coefficient (CPCC) for different sparsity regularization strengths for ON and OFF parasol cells (one retina) and ON and OFF midget cells (one retina). Consensus may be undefined when no subunits are identified as indicated by incomplete stability curves. The dark curves represent the medians (excluding not defined values) of the corresponding population. Stability curves show the expected increase and plateau. (**B**-**D**) Consensus comparison of ON and OFF midget cells. (**B**) Superimposed stability curves (median with 95% confidence intervals shown as shaded regions) of ON and OFF midget cells from (**A**), showing slight but systematic differences. (**C**) Stability curves of ON and OFF midget cells from (**A**) projected onto the first two principal components of all curves from the two populations. (**D**) Inter- and across-type Euclidean distances in the two-dimensional PCA space from (**C**), showing significant differences of distances across versus within cell types. Central line and box represent the median and the interquartile range (IQR; 1st to 3rd quartile), respectively. Whiskers extend to most extreme values within 1.5 IQR, and circles indicate outliers.

Within each type, the shapes of the stability curves exhibit similarities, but this does not equally hold across types. Comparing, for example, the stability curves for ON versus OFF midget cells reveals slight, but systematic differences (**Figure 11B**). The slope of the stability curves for OFF midget cells tends to be somewhat shallower around the bend than for ON midget cells. The difference can be illustrated by analyzing the family of curves with principal component analysis and comparing the shapes of the curves according to the projections on the first two principal components (**Figure 11C**). This shows that curves of cells for the same type tend to group together but diverge across types, despite some overlap of the two populations. It is likely that differences in the levels of noise and in the subunit nonlinearities contribute to these cell-type differences of regularization effects.

More quantitatively, we find that the distances between pairs of curves are significantly larger across cell types than for curves of the same type (**Figure 11D**; two-tailed Wilcoxon rank-sum test, p < 10^−6^ for both ON and OFF midget cells versus across types). This indicates that the stability analysis is similar for cells within a type, but can differ across types, suggesting, along with the tight alignment of the stability curves observed in the salamander dataset (**Figure 7D**), that a suited sparsity regularization parameter can be inferred from a representative subset of cells for each functional cell type. Different types of cells, on the other hand, may require different levels of regularization for optimal performance. The subunit populations in **Figure 10C** are based on cell-type specific regularization parameters, though their differences are small and the effect of these differences rather subtle for the present data.

### Subunits reflect anatomical differences between parasol and midget cell circuits

Midget cells in our analysis yielded on average 5.7±2.3 subunits and parasol cells had 6.3±2.7 subunits (mean ± standard deviation). The sizes of subunits are fairly homogeneous within functional types, but differ between parasol and midget cells (**Figure 12A**; subunit diameters mean ± standard deviation: 60.5±16.7 µm for ON parasol cells, 56.3±10.9 µm for OFF parasol cells, 44.6±12.2 µm for ON midget cells, 38.1±12.9 µm for OFF midget cells, p < 10^−6^ for both ON parasol versus ON midget and OFF parasol versus OFF midget, two-tailed Wilcoxon rank-sum test, data from three marmoset retinas), suggesting distinct presynaptic inputs. Parasol cells in primates receive excitatory inputs from diffuse bipolar cells (Calkins and Sterling 2007; Jacoby et al. 2000; Wässle 1999), whereas midget cells are primarily connected to midget bipolar cells (Jusuf et al. 2006a, 2006b; Kolb and Dekorver 1991).

**Figure 12.**
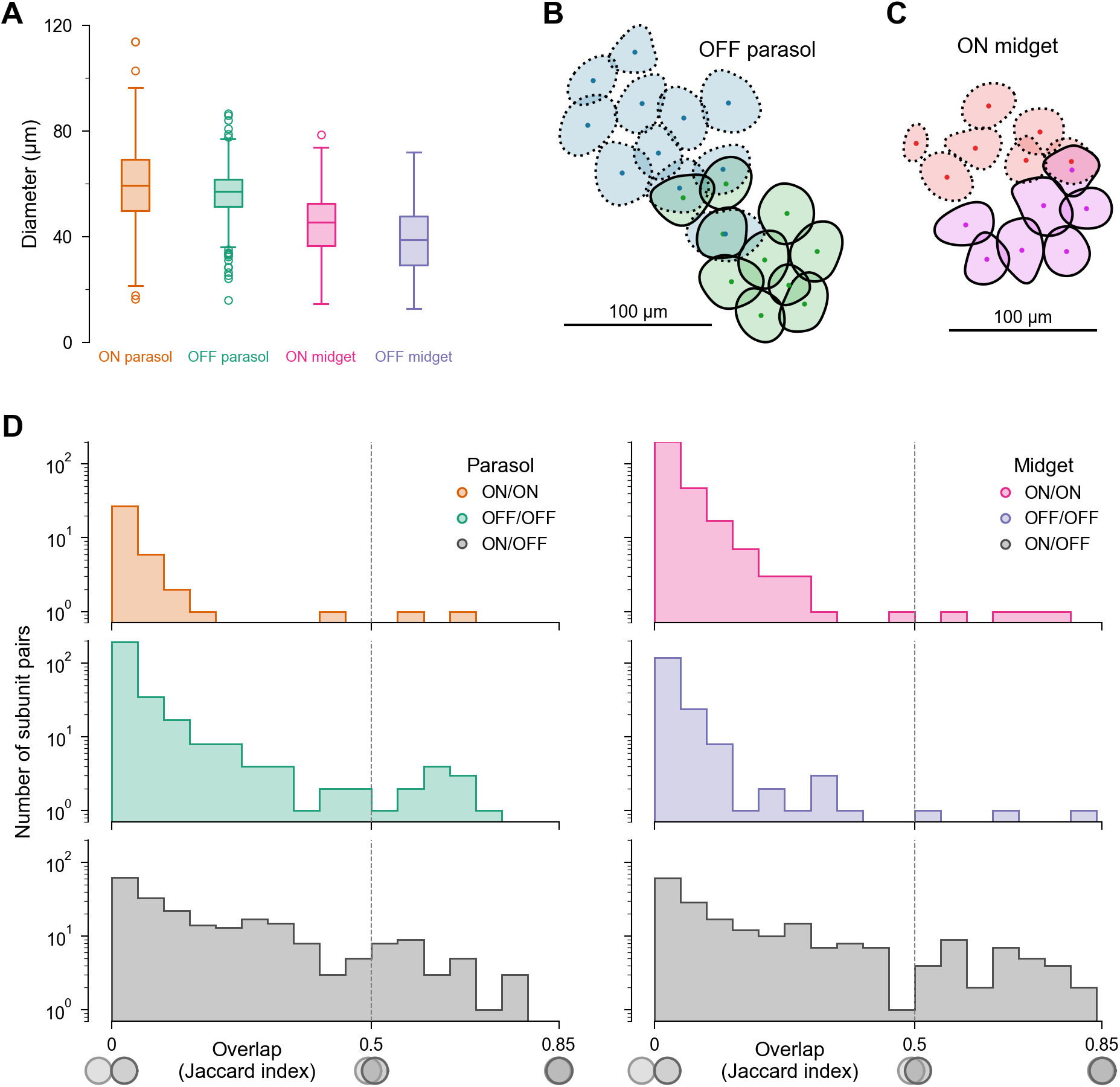
Features of primate retina subunit layouts. (**A**) Subunit diameters of the four major cell types collected from three marmoset retinas. Central line and box represent the median and the interquartile range (IQR; 1st to 3rd quartile), respectively. Whiskers extend to most extreme values within 1.5 IQR, and circles indicate outliers. (**B**-**C**) Examples of the subunits of adjacent parasol (**B**) as well as midget (**C**) cells. Markers represent centers of mass. While the parasol subunits can display substantial overlap from adjacent cells, the midget cell subunit layouts are typically distinct with no or few overlapping subunits. (**D**) Histograms of subunit pair overlap. Overlap between subunit pairs within ON (top), OFF (middle), and across ON and OFF (bottom) parasol and midget cells is measured with the Jaccard index. Only overlap pairs with an index greater than zero are visualized. The dashed line indicates the lower bound of what we consider here as substantial overlap and thus potential shared bipolar cell inputs. Consistent with the examples in (**C**), we observed few overlapping pairs for midget cells, but a considerable amount (**B**) for OFF parasol cells.

For the marmoset retina, there is little information about bipolar cell receptive field sizes, but dendritic trees in the marmoset peripheral retina span around 30 µm for diffuse bipolar cells 15–20 µm for midget bipolar cells in the OFF pathway (Chan et al. 2001). Thus, these relative size differences are consistent with our finding of larger subunits for parasol compared to midget cells, but the absolute values of the recovered subunit sizes are considerably bigger than the dendritic trees. An explanation might be that receptive fields may extend beyond dendritic-tree regions, potentially because photoreceptors may also connect somewhat laterally to bipolar cells or because gap junctions between bipolar cells (Dacey et al. 2000; Kuo et al. 2016; Luo et al. 1999; Manookin et al. 2018) may enlarge receptive fields. For the peripheral macaque retina, for example, diffuse bipolar cells have been reported to have anatomical dendritic tree sizes in the range of 30–50 µm (Boycott and Wässle 1991) and receptive field sizes around 74–114 µm (Dacey et al. 2000), indicating a similar discrepancy between dendritic trees and receptive fields.

### Different ganglion cell types display distinct subunit mosaics

Our results indicate that subunits of ganglion cells belonging to the same functional type arrange in a tiling pattern across the retinal space (**Figure 10C**). Within functional types, one may thus expect that nearby subunits of neighboring ganglion cells either have no or small overlap or overlap to such a high degree that they suggest a common origin, e.g., shared input from the same presynaptic bipolar cell. For OFF parasol cells, we found that the latter was indeed observed, as illustrated by an example of subunits from two cells in **Figure 12B**. Midget ganglion cells also occasionally showed near-identical subunits obtained from two neighboring ganglion cells of the same type (see example in **Figure 12C**), though these were much rarer than for OFF parasol cells. Rather, subunit layouts of neighboring cells complemented each other, jointly tiling space with little overlap. The higher degree of subunit sharing among parasol cells (here in particular OFF parasol cells) compared to midget cells is consistent with anatomical findings. Dendritic trees of midget ganglion cells typically do not overlap, whereas parasol cells do show overlap in the periphery of the primate retina (Dacey 1993; Lee et al. 2010), suggesting the possibility of systematically shared input.

To analyze more quantitatively how the subunits relate to each other, we identified all pairs of subunits from different individual ganglion cells that displayed an overlap and quantified the relative overlap by the Jaccard index (ratio of overlap area and the total area covered by the subunits). The maximum value of this index is unity, which would indicate identical, fully overlapping subunits, whereas values near zero correspond to minimal overlap. **Figure 12D** displays the relative overlap values for parasol cells (left) and midget cells (right). For subunit pairs of ganglion cells from the same type, the vast majority of subunit pairs display a relative overlap below 35%. However, some subunit pairs display particularly strong overlap beyond 50%, especially for pairs obtained from OFF parasol cells. These observations are consistent with a tiling, mosaic-like arrangement of subunits within a functional type, including shared subunits originating from joint input. By contrast, subunits across different ganglion cell types do not show such a mosaic-like arrangement. For parasol versus midget cells, this is already indicated by differences in subunit size (**Figure 12A**). For ON versus OFF parasol cells as well as for ON versus OFF midget cells, subunit pairs display a continuum of overlap values up to about 80% (**Figure 12D** bottom).

### ON- and OFF-type subunit mosaics are aligned

We next compared the large-scale subunit mosaics that emerge from the analysis of ON and OFF ganglion cell populations of corresponding types (midget or parasol) and asked whether these pairs of subunit mosaics display any type of spatial coordination with each other. For example, the layouts of subunits from ON and OFF parasol cells might be aligned with each other, with approximately matching subunit locations between the two mosaics. Alternatively, the mosaics might be anti-aligned with subunit locations of one population lying preferentially in between several subunits of the other population. Or the two mosaics might be independent of each other, displaying no specific coordination. On the level of receptive fields of ganglion cells, a previous study had demonstrated that the mosaics of ON and OFF ganglion cells of corresponding types (parasol ganglion cells in macaque) tend to be anti-aligned, following a prediction from efficient coding theory (Roy et al. 2021). We thus asked whether a corresponding coordination can be observed at the preceding processing layer, as the subunits reflect the organization of the inputs coming to the ganglion cells, which stem from their presynaptic bipolar cells.

For both parasol and midget cells, we selected a recording with particularly dense coverage of receptive fields in order to investigate alignment between ON- and OFF-type mosaics, starting with mosaics of ganglion cell receptive fields (**Figure 13A**). To quantify the level of alignment, we followed the procedure proposed by Roy et al. (2021). We reduced the contour of each receptive field to a single coordinate point, defined by its center-of-mass. We then computed the inter-mosaic coordination energy (IMCE), which describes the pairwise spacing between centroids across mosaics of different cell types (Roy et al. 2021). In this analysis, each heterotypic pair of centroids contributes energy to the IMCE depending on the distance between the two centroids, with smaller distances corresponding to higher energies. Thus, higher values of the IMCE indicate stronger alignment between the mosaics.

**Figure 13.**
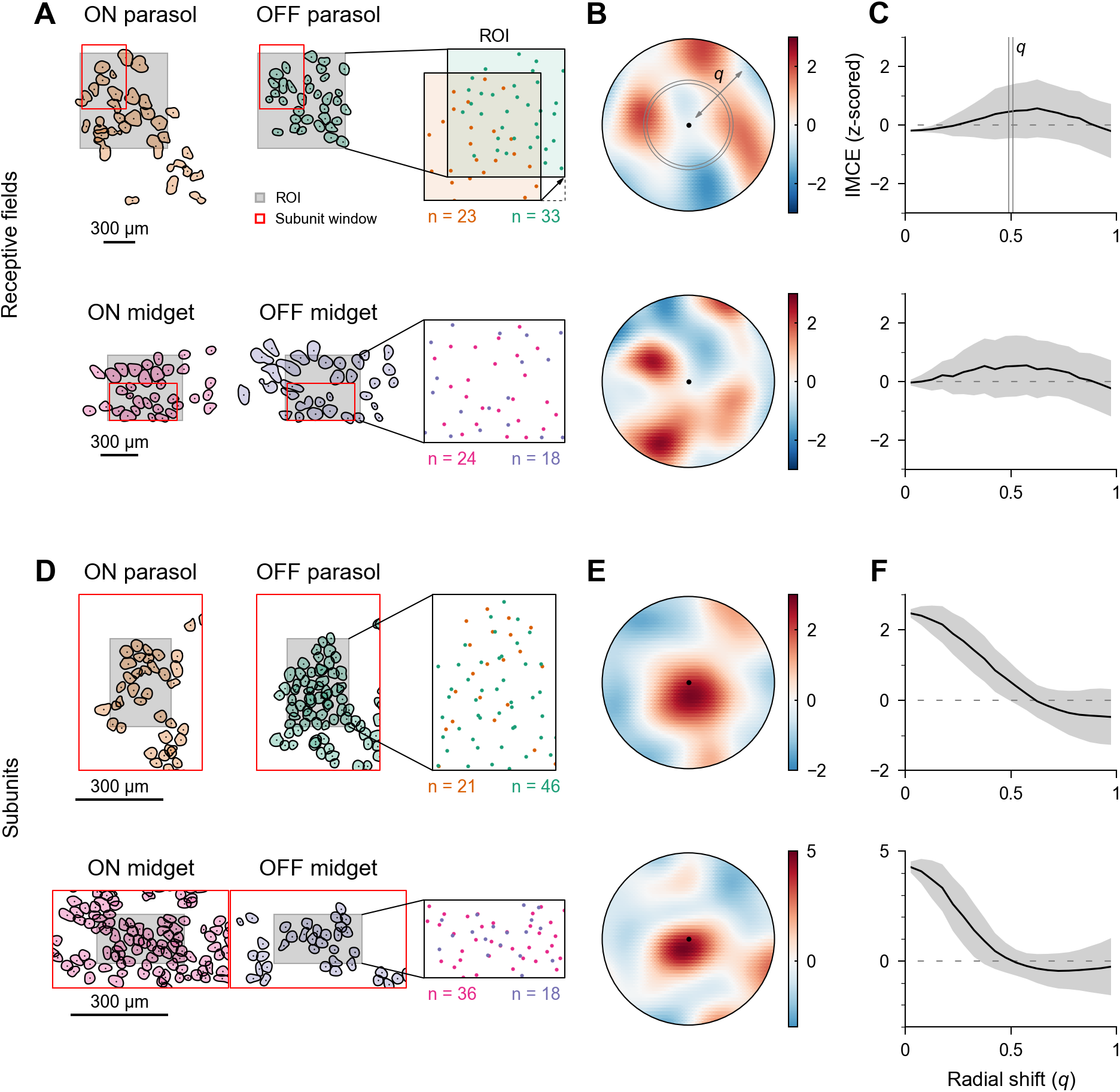
ON- and OFF-type ganglion cell receptive field mosaics are anti-aligned, whereas subunit mosaics show alignment. (**A**) Mosaics of ON- and OFF-type parasol (top) and ON- and OFF-type midget (bottom) ganglion cell receptive field contours, each from one retina, mid-periphery and periphery, respectively. Centers of mass (black markers) inside the region of interest (ROI, shaded area) are considered for analysis (colored markers inside black frame, right). One center-of-mass mosaic is shifted relative to the other, as schematically indicated (right); n = numbers of enclosed centers-of-mass. (**B**) Topographical map visualizing the inter-mosaic coordination energy (IMCE) at different shifts of the ganglion cell receptive field mosaics around the original position in the center. The shift distance *q* is normalized to the median homotypic nearest-neighbor distance of the ON-type mosaic, and data are shown for up to *q*=1, which here corresponds to 89 µm for parasol and 109 µm for midget cells. (**C**) Dependence of the IMCE on radial distance by averaging across angles in (**B**), displaying mean (black) and standard deviation (shaded gray). IMCE topographical map and radial average curves are z-scored, and the shift distance *q* is normalized as explained in (**B**). (**D**-**F**) Same as (**A**-**C**), but for the subunit mosaics of the cell populations analyzed in (**A**-**C**). The shift distance *q* is again normalized to the median homotypic nearest-neighbor distance of the ON-type mosaic, here corresponding to 28 µm for parasol and 29 µm for midget cell subunits. The red rectangles in (**D**) correspond to the red rectangles of the corresponding cell populations in (**A**). The decreasing IMCE curves in (**F**) indicate that corresponding ON and OFF subunit mosaics tend to be aligned.

The analysis consists of observing how the IMCE changes as one mosaic is shifted vertically and horizontally relative to the other (**Figure 13A** right). For aligned mosaics, paired centroids are close to one another and the energy is initially high and decreases with shifting. For anti-aligned mosaics, the energy rises with shifting, as centroids move closer to each other. Finally, for independent mosaics, there is no distinct increase or decrease. We visualize the IMCE for different mosaic offsets as a topographic map (**Figure 13B**). For both parasol and midget ganglion cell receptive field mosaics, the energy increases with the relative shift between the ON and OFF populations, as evident in the radial average of the topographic map (**Figure 13C**). This indicates that the ON- and OFF-type mosaics of the ganglion cell receptive fields are anti-aligned, as previously reported for parasol cells in the macaque retina (Roy et al. 2021).

We then applied the procedure also to the corresponding subunit mosaics (**Figure 13D**) obtained from the same parasol and midget cell populations. Unlike for the ganglion cell receptive fields, the IMCE of the subunit mosaics was found to be largest for zero shift, and a decrease with mosaic shift is apparent for both parasol and midget subunit mosaics (**Figure 13E-F**). This indicates that, in contrast to ganglion cell receptive field mosaics, subunit mosaics obtained from corresponding ON and OFF ganglion cell populations are aligned. Note that mechanistically, such a switch from aligned subunits to anti-aligned receptive fields at the next layer of neuronal processing is not contradictory; ON and OFF ganglion cells may simply pool their signals from (aligned) subunits over distinct, anti-aligned spatial regions.

Note further that both the ganglion cell receptive field mosaics and the subunit mosaics are partially depleted due to missing, non-recorded cells. Although the procedure is robust to subsampling to a certain extent (Roy et al. 2021), we had manually chosen regions of interest for both parasol and midget populations, in which we find few gaps in the tiling for both ON- and OFF-type receptive fields (**Figure 13D** right). The tested portion of the mosaic pairs include around 20–50 subunits each. Performing the analysis with larger regions of interest, defined by the convex hull of centroid pairs excluding outliers, as proposed (Roy et al. 2021), we found that the IMCE remained sharply decreasing with relative shifts of the subunit mosaics, corroborating the alignment of the ON- and OFF-type subunits.

## DISCUSSION

We propose a method for subunit identification based on spike-triggered non-negative matrix factorization (STNMF), which finds subunits within a receptive field from recorded spike-train data in a matter of seconds of computation time. The approach enables, for example, large-scale analyses of the primate retina, recovering the nonlinear subunits of the four major ganglion cell types: ON and OFF parasol and midget cells. This allows in-depth cell-type-specific analyses of subunits on a cell population level, comparison to type-specific anatomical properties, and new insights about the spatial coordination of ON- and OFF-pathway signal transmission.

### Recovered subunits resemble bipolar cell receptive fields in the primate retina

We applied the method to recorded spike trains from marmoset retinas and analyzed their subunit properties. Subunits are generally thought to results from nonlinear bipolar cell input to the ganglion cells. For example, primate parasol cell subunits have been suggested to correspond to diffuse bipolar cells (Crook et al. 2008, 2014). We found that parasol cells yielded larger subunits than midget cells, though also somewhat larger than dendritic trees of diffuse bipolar cells in the peripheral marmoset retina (Chan et al. 2001). Note, though, that the bipolar cell inputs do not occur in isolation, but are shaped and modulated by interactions with amacrine cells (Jacoby et al. 1996; Kolb and Dekorver 1991) and gap junctions between individual bipolar cells (Appleby and Manookin 2020; Jacoby et al. 2000; Manookin et al. 2018). Together, for parasol cells, our findings concerning subunit density, uniform tiling, and potential sharing of subunits between pairs of parasol cells are at least consistent with the notion that the parasol subunits may correspond to individual bipolar cell inputs.

Midget ganglion cells appear to receive input from around eight (mid-periphery) up to 13 midget bipolar cells (periphery) in the marmoset retina (Jusuf et al. 2006b). We typically found fewer subunits per midget ganglion cell. Yet, these subunits of midget cells form a mosaic at the population level and subunit sizes seem consistent with what one might expect for receptive fields of individual bipolar cells. It is possible that the discrepancy in the numbers of subunits and connected bipolar cells reflects that some subunits do not correspond to individual bipolar cells or that some subunits remain undetected. Alternatively, some anatomically connected bipolar cells might not have a strong functional connection to the postsynaptic midget cell under the applied spatiotemporal white-noise stimulus.

Furthermore, some ganglion cells, in particular parasol ganglion cells, can be connected to multiple types of bipolar cells (Jacoby et al. 2000; Tsukamoto and Omi 2016). The identified mosaics of tiling subunits in our analysis, however, rather suggest correspondence to a single type of bipolar cells. Although STNMF can in principle identify subunits with considerable overlap (Liu et al. 2017), practically there will be a bias towards the dominant subunits, that is, those that provide the stronger drive or the more nonlinear input. Similarly, nonlinearities coming from earlier processing stages, as observed in photoreceptor signals or bipolar-cell input (Angueyra et al. 2022; Endeman and Kamermans 2010; Grabner et al. 2023; Schreyer and Gollisch 2021), could in principle affect identified subunits, but may not be effective if the later, e.g., bipolar-cell-mediated, nonlinearities dominate.

### Subunits arrange in a uniform mosaic by cell type

Our subunit analysis with STNMF revealed distinct mosaics of subunits for ON and OFF parasol as well as ON and OFF midget cells in the marmoset retina, in line with ganglion cell types stratifying in different levels of the inner plexiform layer (Watanabe and Rodieck 1989) and receiving input from distinct bipolar cell types (Masland 2001; Wässle 2004). The observed uniformity of the subunit mosaics matches reports of constant bipolar-cell input density across ganglion cell dendritic trees and across retinal space in the marmoset (Eriköz et al. 2008).

There is recent evidence that the mosaics of ON and OFF parasol ganglion cell receptive fields in the macaque retina are spatially anti-aligned (Roy et al. 2021). Models of efficient coding indicate that anti-alignment is the optimal arrangement between populations of ON and OFF cells in a high-noise/high-threshold coding regime, in which ganglion cells likely operate, as it allows nearby ON and OFF cells to encode non-redundant information (Jun et al. 2021). Increased resolution is an additional benefit of anti-alignment, and it may have advantages for down-stream convergence (Roy et al. 2021). Our present analysis confirmed the anti-alignment of ganglion cell receptive fields also in the marmoset retina. By contrast, the same analysis applied to the identified subunit mosaics indicated alignment rather than anti-alignment between ON- and OFF-type subunit mosaics for midget as well as for parasol cells. Under the assumption that the subunits indeed correspond to receptive fields of bipolar cells, there thus seems to be a switch from aligned to anti-aligned spatial coordination when going from the bipolar cell to the ganglion cell layer.

From a functional perspective, a reason for this switch could be that the conditions that make anti-alignment efficient for ganglion cell receptive fields do not hold at the level of bipolar cells. Owing to the smaller receptive fields of bipolar cells compared to ganglion cells, anti-alignment may offer little additional benefit in spatial resolution. Furthermore, whether alignment or anti-alignment is more efficient in terms of information transmission depends on the noise regime in the system, with alignment becoming optimal as lower noise levels are considered (Jun et al. 2021). Noise at the bipolar cell level, where signals are carried by graded potentials rather than stochastic spikes, might effectively be lower than in ganglion cells, which are affected by noise in the bipolar-to-ganglion cell synaptic transmission (Borghuis et al. 2009; Freed and Liang 2014) and by the spike-generation process (Berry et al. 1997), and this could favor alignment at the bipolar cell stage. Yet, a primary noise source in the retina occurs at the level of photoreceptors (Ala-Laurila et al. 2011; Angueyra and Rieke 2013) and thus also affects bipolar cell signals, and the effects of these noise sources on information transmission also depend on whether and how the signals from ON and OFF bipolar cells might be jointly read out.

Mechanistically, the alignment of ON and OFF subunits might be aided by the fact that ON and OFF bipolar cells receive their input from the same presynaptic cells, the cone photoreceptors under our photopic light conditions. In the macaque retina, for example, subunits of OFF midget ganglion cells have been shown to often contain just a single cone location (Freeman et al. 2015), which may lead to naturally aligned subunits if the same cone forms the basis of a subunit for both an OFF and an ON midget cell. In the marmoset retina, however, cones are several times more densely packed than for macaques, with densities reported around 10,000 to 20,000 cones per mm^2^ in the periphery (Troilo et al. 1993; Wilder et al. 1996), suggesting that the subunits identified here cover a dozen or more cones. Thus, the recovered subunits of marmoset ganglion cells cannot be directly related to the underlying spatial pattern of cones, and the subunit alignment can therefore not easily be traced back to joint input from individual cones.

### Accelerated Fast HALS as a fast and versatile method for STNMF

The large-scale analyses were made possible by the improved methodology of different aspects in the STNMF approach, including purpose-built initialization, a tailored NMF algorithm, and adjusted consensus methods for hyperparameter selection. The obtained reduction in computational time is useful in several ways. First, it considerably facilitates analyses of large data sets. Second, it allows probing and optimizing various parameters, such as regularization strength, but also the size of the considered spatial or temporal window or the number of modules. And third, it lets a user quickly and easily explore the method and build intuition regarding its applicability to a given dataset and regarding the limits and possibilities of the various choices to make for the analysis. Although some of these points may also be realized by improvements in computing hardware or parallelization, optimization in the multidimensional space of hyperparameters is best served by combining fast hardware with efficient algorithms. And the availability of an easy entry point, yielding rapid initial results without the need for hardware optimization, may be crucial for whether the method is applied or not.

Solving a non-negative least squares problem (NNLS) sits at the core of NMF. To solve it accurately, typically, costly algorithms are employed iteratively (Kim and Park 2011). These include multiplicative updates (Lee & Seung, 1999), the projected quasi-Newton method (D. Kim et al., 2007), the projected gradient method (Lin, 2007), non-negative quadratic programming (Zdunek & Cichocki, 2008), and the active-set method (Lawson & Hanson, 1974). The algorithms strive to provide exact solutions at each iteration by taking into account all data at once. Here, we avoid the high computational demand with hierarchical alternating least squares (HALS) by solving smaller, non-restricted least squares problems followed by setting all negative values to zero (Cichocki et al. 2007; Ho 2008; Li and Zhang 2009). While Fast HALS (Cichocki and Phan 2009) simplifies computational operations for speed, A-HALS (Gillis and Glineur 2012) refines the coarse approximations of HALS by performing multiple cycles of updates within one NMF iteration. The intended purpose of A-HALS is to accelerate the convergence by reducing the number of necessary iterations, which lends it its name. However, by incorporating sparsity regularization, we make use of another advantage of this procedure. Additional constraints like sparsity regularization are typically integrated into the equations of the iterative update (Kim and Park 2007; Liu et al. 2017). As we instead perform it multiple times, that is, at each A-HALS cycle within one NMF iteration, the continuously applied sparsity constraint refines the subsequent repetitions by giving more weight to prominent structures and progressively suppressing the effect of uncorrelated noise. Consequently, the result after one full iteration offers relatively high accuracy and sparsity. Our AF-HALS method with sparsity regularization may prove to be a valuable tool not only for the estimation of subunits, but also for other applications of NMF.

### Sparsity regularization as a crucial ingredient of STNMF

In the field of machine learning, sparsity regularization has become an important tool to combat overfitting and has earned the role of an enhancement strategy to allow generalization when data is limited. In many applications of signal processing and learning representations, however, sparsity has proved to be an even more essential and critical player (Candès et al. 2006; d’Aspremont et al. 2007; Donoho 2006; Hoyer 2004). Sparsity in NMF offers interpretable solutions (d’Aspremont et al. 2007) and inherently reduces the impact of noise by decreasing the effects of uncorrelated patterns in inputs (Donoho 1995; Elad 2006; Hyvärinen 1999). Aside from these benefits, sparsity is an integral part of STNMF. It replaces the dropped non-negativity constraint of one factor matrix in semi-NMF (Hoyer 2002, 2004; Kim and Park 2007, 2008), as otherwise the parts-based reconstruction can collapse (Ding et al. 2010). In the context of STNMF, sparsity is thus a vital component for producing localized subunits.

Beyond simple *ℓ*1-norm-based sparsity, there have been other efforts towards regularization techniques that favor localized solutions in the fields of NMF and clustering such as decomposing widefield calcium imaging signals into localized functional brain regions (Saxena et al. 2020). The prior definition of the localized regions based on a brain atlas in this previous study, however, does not easily translate to our problem of identifying subunits of unknown number, locations, and shapes. A different approach to enforce localized solutions is to apply a locally normalized *ℓ*1-regularization, which was introduced with spike-triggered clustering (Shah et al. 2020) to instill spatial information into the *ℓ*1-regularization in subunit estimation. The penalty of sparsity regularization at a given location of a subunit was adjusted based on the spatially neighboring values. For our application, the forced locality is not directly applicable, as it makes a module either contain a localized pattern or collapse to no non-zero values. Noise in the input cannot easily escape to excess modules, and the number of included modules would have to be precisely determined instead of using an upper bound on the number of expected subunits as is done here. An alternative approach to regularization with localized structures comes from using basis functions to parameterize stimulus filters. For example, using spline-based parametrizations can help estimate receptive fields and has also been combined with subunit-based model fits (Huang et al. 2021).

### Parameter selection by consensus analysis

To find a suitable amount of sparsity regularization, we investigated how robust the decomposition is with respect to varying initialization. To measure how stable the decomposition is, we here adjusted the consensus procedure proposed by Brunet et al. (2004) to be applied to the problem of identifying regularization parameters. An alternative to using consensus analysis is L-curve fitting. The L-curve technique (Hansen 1999, 2001) aids in determining a suitable parameter for a regularized least squares problem. The method inspects the relation between the residuals and the norm of the solution. With too much regularization, the data will not be properly accounted for, and with too little regularization, the fit will be good but dominated by undesired effects. Both terms are visualized against each other in logarithmic space for a range of regularization values, exhibiting a curve shaped like the letter “L”. Determining a corner point can then be used to obtain a suitable trade-off between the two terms for the least squares optimization and the regularization penalty. An advantage over our proposed consensus analysis is the conceptual simplicity and that it does not require multiple repetitions of NMF. It has been applied to regularized NMF with HALS previously, although for a different type of regularization (Cichocki and Phan 2009). Nevertheless, when applied to STNMF, we did not find a sharp corner with the data available, and also the dedicated method of L-corner fitting (Hansen 1994) did not succeed. We therefore turned towards the more involved consensus analysis, which we found to provide a good alternative.

### NNSVD-LRC offers reliable initialization

We demonstrate that a single run of STNMF with SVD-based initialization can replace the selection of the most suitable solution among 100 randomly initialized runs. Specifically, using NNSVD-LRC is advantageous, as it makes NMF algorithms converge faster to a stationary point due to low error and increased sparsity in the initialization (Atif et al. 2019). Notably, on the one hand, techniques based on SVD do not guarantee NMF to find the global optimum (Atif et al. 2019). On the other hand, however, seeking the global optimum in network structures bears the risk of overfitting (Choromanska et al. 2015). Furthermore, the alternative of selecting a most suitable among many randomly initialized solutions raises challenges because a measure of suitability is not easily defined and a manual selection of a best solution or a particular criterion is subject to human bias. To alleviate these problems, an adapted consensus clustering procedure (Zhou et al. 2020) poses a viable alternative for initialization. Instead of choosing the best decompositions among 100 randomly initialized NMF runs, the resulting modules are concatenated to serve as the input to another NMF run that yields a final set of modules. We found that decompositions initialized with NNSVD-LRC are comparable with solutions of that approach while requiring only one run of NMF.

### Limitations

Our results show that STNMF has strong potential to identify subunits of receptive fields under certain conditions, but there are also important limitations. Firstly, the method poses constraints on the stimulus with which the spiking data is obtained. To infer subunits from pixel correlations in the spiking response, a white-noise stimulus with uncorrelated pixel contrast across space is necessary. This is typically not a particularly strong stimulus for activating retinal ganglion cells, which means that adequate time needs to be allocated to this specific stimulus during the experimental recording to gather sufficient numbers of spikes. Here, we were able to recover subunits successfully from less than one hour of recording time. Whether the amount of data is sufficient to reliably identify subunits can be checked by analyses that use only a subset of spikes, as done in **Figure 9**. This analysis also shows that the number of extracted subunits does not per se increase with the amount of data, at least when sufficient numbers of spikes are available.

It is also worth mentioning that the need to use stimuli with fine spatial resolution, such as white noise, is intrinsic to the nature of the problem of localizing subunits, independent of the analysis method. Natural stimuli, for example, may be limited for localizing subunits by their relative lack of power at high spatial frequencies, where the exact locations of subunits actually matter. An alternative stimulation approach, which combines high spatial resolution with good activation of ganglion cells, is to use flashed or randomly flickering gratings (Karamanlis et al. 2025), though inferring subunits then requires fitting an adequately parameterized computational model.

Another caveat is the pixel size of the stimulus. Although STNMF makes few assumptions on the data, the stimulus design must provide stimulus pixels of suitable size to fit bipolar cell receptive fields, whose spatial scales can vary with species and with location on the retina. To aid stimulus design, estimates of dendritic-tree size or measured receptive fields in the literature can be consulted, or relevant spatial scales of nonlinear integration can be estimated from responses to standard reversing-grating stimuli at varying spatial frequency.

A third limitation is that we here used a single temporal filter for a given ganglion cell, derived from the cell’s STA, to integrate out the temporal stimulus dimension. This assumes that a ganglion cell’s subunits all have at least approximately the same temporal filter, which may be violated if, for example, subunits are formed by different types of bipolar cells. For the present application, the reason for relying on a single temporal filter is twofold. First, we have shown in previous work (Liu et al. 2017) that this approach also works when the temporal filters of different subunits are not identical, but overlap to at least some extent because this will let the effective spike-triggered stimuli still carry correlations that reflect the subunits. Second, uniform temporal filters seem a good approximation for the subunits of individual ganglion cells because inputs are likely dominated by a single type of bipolar cells, even if others also contribute activation. Subunits then likely reflect the dominant input type. For a future extension of the method, it would in principle be possible to allow subunits to have their own temporal profile, though this would lead to a considerable increase in the size of the matrices of the NMF problem and thereby raise challenges regarding required data and computational time. An alternative might be to include a low-rank approximation of the spatiotemporal modules by enforcing a space-by-time separation with a small number of temporal kernels, as has been previously used in NMF-based analyses of neuronal population activity (Onken et al. 2016).

Finally, STNMF as used here so far cannot produce subunits with both positive and negative components. Although the unconstrained sign of the STNMF weights allows the recovery of subunit layouts containing simultaneously subunits with purely positive as well as with purely negative entries (the latter recovered with inverted sign, but associated with negative weights), mixed structures, such as subunits with an antagonistic surround, are not consistent with the non-negativity constraint. One might speculate that antagonistic structures could be recovered as separate, additional subunits if their effects are strong enough, but we have not observed such cases in our datasets, consistent with the fact that surround components are often weak under spatiotemporal white-noise stimulation (Wienbar and Schwartz 2018). The non-negativity constraint also seems to preclude application of the method to many systems where subunits might be expected to have both On-type and Off-type components, such as primary visual cortex (Rust et al. 2005). However, workarounds may come from using stimulus parametrizations that allow composing subunits as non-negative superpositions of basis functions, which themselves may contain positive and negative components, such as wavelets. STNMF would then work with the weights of the wavelets at different positions rather than directly with the pixel contrast values.

In fact, an insightful analysis can already be obtained by including separate stimulus components for capturing On- and Off-type sensitivity, respectively, within the same subunit. This is illustrated by the analysis of a simple model for a complex cell in primary visual cortex (**Figure 14A**). The “trick” here is to use two (normally distributed) random numbers per stimulus pixel and interpreting one as a positive-contrast component and the other as a negative-contrast component (so that positive values designate negative contrast and vice versa for this component). The actual stimulus is then given by the difference of the two stimulus components, but the subsequent analysis treats the two sets of random numbers as separate stimulus components (**Figure 14B**). This effectively doubles the number of pixels in the analysis, but lets each subunit “choose” whether to incorporate the positive-contrast component or the negative-contrast component for each pixel. The STA (**Figure 14C**) reflects the two components by showing the receptive field and its inverted version in the two component regions. The standard STNMF analysis (**Figure 14D**) then recovers the subunits with their On-center in the positive-component part (top half of each displayed module) and the Off-sidebands in the negative-component part (bottom half of each module). In addition, each subunit is detected a second time, now with a sign-flip because the On-center appears in the negative-component part. This occurs because stimuli with negative numbers in the negative-component part activate the On-center of the subunits, and STNMF can reconstruct these stimulus patterns by combining non-negative module entries in the negative-component part with negative entries in the weight matrix. Yet, despite the artificial doubling of the detected subunits, the STNMF results clearly indicate three spatially-offset On-center/Off-sideband subunits, as were used in the simulation. Furthermore, the subunit recovery here succeeded despite the overlap between the On-centers of the Off-sidebands of the neighboring subunits.

**Figure 14.**
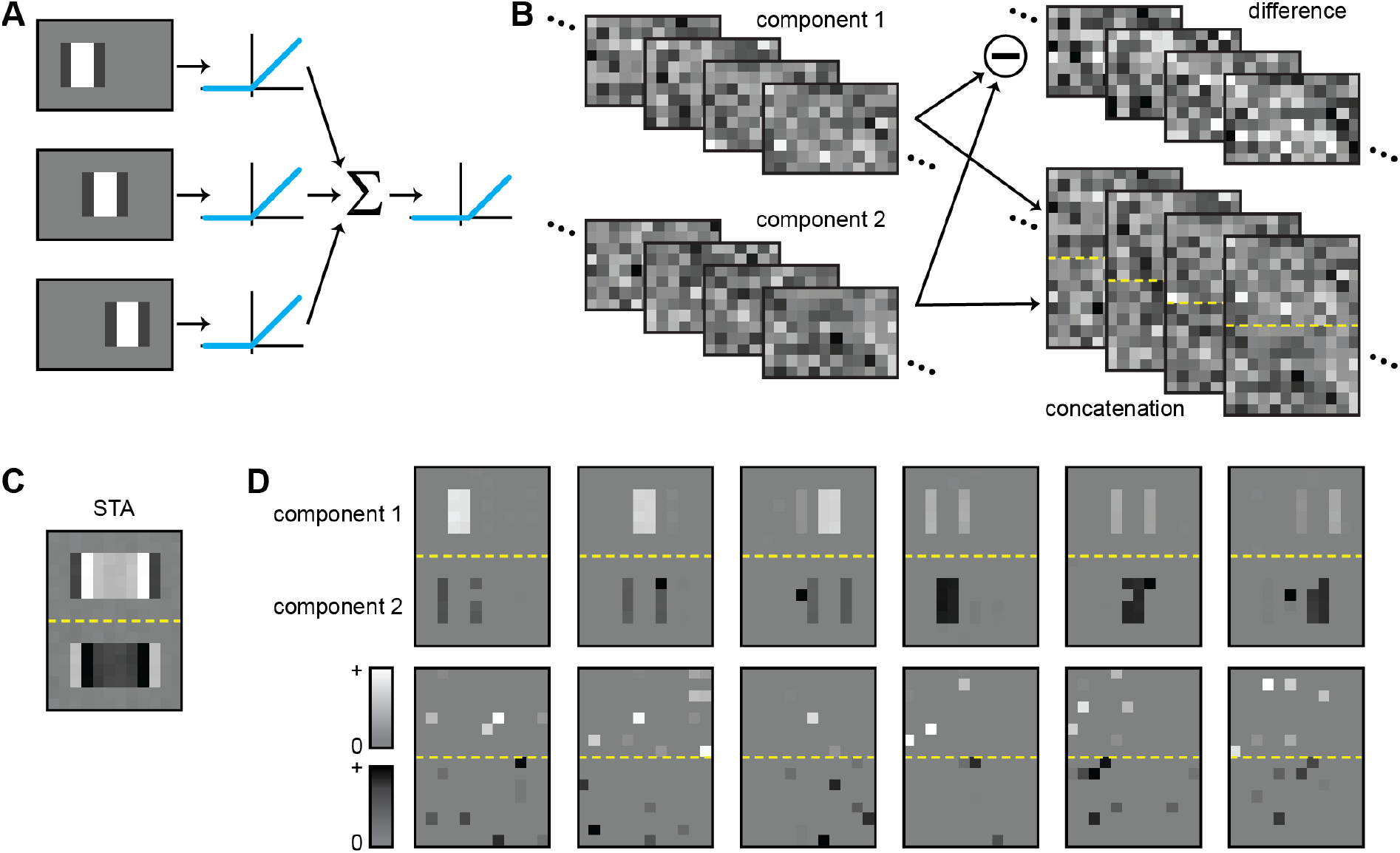
Identification of subunits for a simulated complex cell of primary visual cortex. (**A**) Schematic of the simulation. Three spatial subunits, each with an On-center and Off-sidebands, were used to filter the stimulus (here purely spatial with no temporal structure). The subunit outputs were thresholded at zero and summed, before another thresholding nonlinearity was applied with slightly positive threshold. Based on the resulting activation, a Bernoulli process was used to determine which of the input stimulus patterns generated spikes. (**B**) The stimulus was composed of two components (left), each corresponding to a sequence of spatial Gaussian white-noise patterns. The difference of the two components (top right) was used for activating the subunits, the concatenated patterns (bottom right; dashed yellow line denoting the separation between the regions of components 1 and 2) for the subsequent analysis (STA and STNMF). (**C**) The spike-triggered average (STA) reveals the actual receptive field, corresponding to a superposition of the three subunits, in the top part and an inverted version resulting from the use of stimulus component 2. (**D**) STNMF reveals subunits with On-center components in the region corresponding to stimulus component 1 and Off-sidebands in the region of stimulus component 2 (top row, first 3 modules), as well as inverted versions of these subunits (top row, last 3 subunits). Modules are here plotted with different grayscale schemes in the two regions, using gray (for zero) to white (for positive components) in the top part, and gray to black in the bottom part, to signify that positive values correspond to positive contrast in the top half and to negative contrast in the bottom half. The present analysis was based on 1,000,000 input patterns, leading to about 170,000 spikes, and on STNMF with 12 modules and a regularization coefficient of 1.8.

### Comparison to other methods of subunit inference

Popular computational methods of subunit inference in recent years have been linear-nonlinear cascade models (Maheswaranathan et al. 2018; Real et al. 2017), approaches of convolutional neural network models (Maheswaranathan et al. 2023; McIntosh et al. 2016; Tanaka et al. 2019), and methods of statistical inference (Liu et al. 2017; Shah et al. 2020). Cascade models, for example, can include subunits as a layer of parallel linear stimulus filters and be trained in a supervised manner on the relation between stimulus input and spiking response. This approach was successfully applied to data from retinal ganglion cells of salamanders (Maheswaranathan et al. 2018; Real et al. 2017), using a stimulus with spatial variations restricted to one dimension to facilitate parameter fitting.

Convolutional neural networks are similarly trained end-to-end. The models are often optimized on data from a population of neurons and provide convolutional filters of subunit types rather than individual subunits for each neuron. Training can be done with natural images instead of artificial stimuli like white noise or flickering bars. For models trained with ganglion cell data from the salamander retina, obtained convolutional filters shared properties with actual bipolar and amacrine cells (Maheswaranathan et al. 2023; Tanaka et al. 2019), although it remains unclear to what extent the model architecture resembles the actual neural circuitry. Spike-triggered clustering (Shah et al. 2020), on the other hand, recovers subunits on stimulus correlations much like STNMF. The obtained models were capable of predicting spiking responses to natural scenes for primate retinal ganglion cells, but subunit number had to be determined through cross-validation, unlike for STNMF.

### Beyond retinal subunits

Nonlinear signal integration is a ubiquitous feature of neural systems and of sensory signal processing in particular. Analyses based on receptive field subunits are thus applicable to many other sensory areas and have been applied in primary visual cortex (Almasi et al. 2020; Bartsch et al. 2022; Lehky et al. 1992; Vintch et al. 2015), in medial superior temporal visual cortex (Beyeler et al. 2016; Mineault et al. 2012), and in the auditory system (Ahrens et al. 2008; Harper et al. 2016; Keshishian et al. 2020; McFarland et al. 2013). The methodology presented in the present work can also be applied to these systems, given sufficiently long recordings with white-noise-like stimuli and appropriate stimulus parametrizations (c.f., **Figure 14**). In the case of the auditory system, for example, subunits might then correspond to filters in spectro-temporal space, whereas subunits in higher, motion-sensitive visual cortical areas may be partly defined by their direction and speed tuning. More generally, employing STNMF in these systems should aid in developing nonlinear stimulus–response models that link the functional circuitry to the performed neural computations.

In addition to sensory systems, the implementation of NMF developed here together with the presented techniques for controlling hyperparameters and initialization should be of interest for analyzing neuronal population activity. Uncovering hidden structure and dynamics in the joint activity of large ensembles of neurons is seen as an important part of understanding neural computations and information flow in the brain, and a common approach is to identify latent variables of the population activity via dimensional-reduction techniques (Cunningham and Yu 2014; Gallego et al. 2017; Gao and Ganguli 2015; Koh et al. 2022). NMF provides an appealing approach to this challenge, as one may hope that the non-negativity constraint helps infer latent variables—the equivalent of the subunits of the ganglion cell receptive fields—with interpretable structure (Carbonero et al. 2024; Devarajan 2008; Nagayama et al. 2022; Onken et al. 2016). The computational speed achieved by the combination of fast and accelerated HALS as demonstrated in this work should be advantageous for applications of NMF to neural population data, in particular in order to control regularization and other hyperparameters.

## ACKNOWLEDGMENTS

This work was supported by the European Research Council (ERC) under the European Union’s Horizon 2020 research and innovation programme (grant agreement number 724822), by the Else Kröner Fresenius Foundation through the Else Kröner Fresenius Center for Optogenetic Therapies, and by the Deutsche Forschungsgemeinschaft (DFG, German Research Foundation) – project numbers 432680300 (SFB 1456, project B05), 528760423 (SFB 1690/1, project B05), and 515774656.

## AUTHOR CONTRIBUTIONS

SJZ developed and implemented the AF-HALS-based STNMF method, collected and curated the marmoset retina data, analyzed the data, and prepared figures and the initial draft of the manuscript. DK, MHK, HMS, SS, VR, SK, MM, and DAP all contributed to experiments and data collection. SJZ, DK, MHK, HMS, and TG supplied methodology. TG supervised the project and contributed to analyzing the data, performing model simulations, preparing figures, and writing the manuscript. All authors helped revise the manuscript.

## DATA AND SOFTWARE AVAILABILITY

The salamander spike trains analyzed in this work come from a publicly accessible repository (https://gin.g-node.org/gollischlab/Liu_etal_2017_RGC_spiketrains_for_STNMF; DOI: 10.12751/g-node.62b65b). We have made the marmoset spike trains from the present study publicly available in this repository: https://gin.g-node.org/gollischlab/Zapp_Gollisch_2023_Marmoset_RGC_spiketrains_under_spatiotemporal_white_noise (DOI: 10.12751/g-node.zpj6rc). The implemented software for STNMF analysis is provided as a Python package. It includes the NMF algorithm and initialization procedures optimized with NumPy for multi-threaded CPU computation. Tools for visualization are also available as well as routines for the consensus analysis and custom extensions via callback functions. The software, along with an extensive documentation and example code for running the analysis, is available in a GitHub repository: https://github.com/gollischlab/STNMF_with_AFHALS.

## METHODS

### Ethics statement

All experimental procedures were performed in strict accordance with national and institutional guidelines. Recordings from marmoset retinas were performed on tissue obtained from animals used by other researchers, as approved by the institutional animal care committee of the German Primate Center and by the responsible regional government office (Niedersächsisches Landesamt für Verbraucherschutz und Lebensmittelsicherheit, permit number 33.19-42502-04-17/2496).

### Electrophysiology

The salamander data stem from a publicly available repository (Gollisch and Liu 2018) and had been described and analyzed in a previous publication (Liu et al. 2017). Here, we analyzed spike trains of the first retina recording in the repository (“cell_data_01_NC.mat”). The spike trains had been recorded from ganglion cells in isolated axolotl salamander (*Ambystoma mexicanum*) retinas, mounted on 252-channel planar multi-electrode arrays (MEAs) and provided with oxygenated Ringer’s solution at a constant temperature of around 22°C.

Data from common marmoset (*Callithrix jacchus*) were recorded from retinas of three animals of different sex and age (female 2.3 years, male 18.1 years, and male 13.2 years). The eyes were removed between three and twelve minutes after time of death. Left and right eye were labeled and dark-adapted for at least one hour. The retina was isolated and cut into pieces, classified at time of dissection by their quadrant (nasal, superior, and dorsal) and eccentricity (central, mid-periphery, and periphery). The temporal quadrant was not considered due to the optic disc and the high density of cell bodies and axons at the fovea. Dissection was performed under infrared illumination on a stereo-microscope equipped with night-vision goggles. Individual retina pieces were removed carefully from the sclera and choroid and placed on MEAs (MultiChannel Systems, Reutlingen, Germany). Pieces not recorded immediately were stored in darkness in oxygenated (95% O2 and 5% CO2) Ames’ solution, supplemented with 6 mM D-glucose, and buffered with 22 mM NaHCO3 to maintain a pH of 7.4. The data analyzed in this work were recorded from three pieces (one mid-periphery, two periphery), using either a conventional 60-channel MEA (10 µm electrode diameter, 100 µm electrode distance; one recording) or a 60-channel perforated MEA (30 µm electrode diameter, 100 µm electrode distance; two recordings), containing holes between the electrodes in order to keep the retina piece in place by slight suction from below. On the conventional MEA, the piece was held in place by coating the MEA with poly-D-lysine (Millipore, 1 mg/mL), applied to the array for one to two hours and then rinsed off prior to mounting.

During recording, the retina pieces were perfused with oxygenated Ames’ solution and kept at a constant temperature of around 32–34°C, using an in-line heater and, in the case of the conventional MEA, additionally a heating plate below the array. Bandpass-filtered (300 Hz to 5 kHz) voltage signals from each electrode were sampled at 25 kHz. Spikes were extracted using Kilosort (Pachitariu et al. 2016) adjusted to accommodate for MEA data (https://github.com/dimokaramanlis/KiloSortMEA). The automated extraction was followed by visual inspection and curation using Phy2 (https://github.com/cortex-lab/phy). Clusters that displayed good separation from other clusters as well as a uniform spike waveform and a clear refractory period were considered as representing individual ganglion cells.

### Visual stimulation

Spiking activity was recorded in response to visual stimulation. Stimuli were projected onto the photoreceptor side of the retina using a gamma-corrected white-light OLED projector (eMagin, USA) at 800 × 600 square pixels with a refresh rate of 60 Hz (salamander and two marmoset experiments) or 85 Hz (one marmoset experiment). The pixel size (width and height) translated to 2.5 µm or 7.5 µm on the retina, depending on the configuration. The projector was controlled by a custom-built software based on C++ and OpenGL to generate the stimulus frames. The spatiotemporal white-noise stimulus was presented with a pixel size of 15 µm on the retina (30 µm for salamander) and temporally updated at 30 Hz (salamander), 20 Hz, or 21.25 Hz (marmoset). The light intensity of each stimulus pixel was drawn independently from a binary distribution with 100% contrast and a mean light level of about 2.5 mW/m^2^ (salamander), 3.33 mW/m^2^, or 4.89 mW/m^2^ (marmoset). For analysis, the stimulus was described by its contrast values of +1 (bright) or –1 (dark) for each stimulus pixel. The white-noise stimulus was presented in (non-repeating) sections of 50–180 seconds (depending on the experiment), which were periodically interleaved with a presentation of the same, fixed white-noise sequence (10–31 seconds), intended as held-out data for model validation. The data from the fixed white-noise sequence were not considered in this work. The recording durations of the marmoset experiments ranged between 43 minutes and three and a half hours of non-repeated stimuli (3:28, 0:43, and 1:42 hours).

### Receptive field analysis

The receptive field of a given ganglion cell was estimated using reverse correlation to obtain the spike-triggered average (STA; Chichilnisky 2001). The stimulus frames within the 700 ms time window preceding a spike were collected and averaged across all spikes. The STA summarizes the spatiotemporal dynamics of the receptive field. To describe the temporal filter of the STA, we selected the pixel that exhibits the maximum absolute value across all STA frames. We defined the temporal filter as the value of that pixel across time, normalized to unit Euclidean norm. The spatial profile was defined as the stimulus frame of the STA that contained the highest absolute pixel value.

The receptive field diameter was estimated by fitting a two-dimensional Gaussian function to the spatial profile of the STA. The effective receptive field diameter 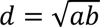 was obtained from the major axis *a* and minor axis *b* of the ellipse at two standard deviations.

For inspection of subunit shape and tiling, contour outlines of the receptive field and subunits were produced. First, the image of each spatial filter was up-scaled by a factor of eight with nearest neighbor sampling and then smoothed with a circular Gaussian filter with a standard deviation appropriate to the spatial size (1.2 original stimulus pixels for receptive fields and 0.5 stimulus pixels for subunits). The contour was determined with the function “find_contours” of the Python module “skimage.measure” at 47.5% of the maximum value of the smoothed filter, equivalent to the 1.22-sigma contour of a circular Gaussian fit. To counter possible expansion effects by the Gaussian smoothing, two-dimensional Gaussians were fit to the smooth and non-smooth filter image. The ratio in their effective diameters was then used to correct and rescale the contour by scaling the contour-level sigma. Contour islands and holes were excluded by selecting the positive closed contour of largest area. Contours were used for visualization of the subunit layouts and mosaics, as well as for determining the relative overlap of two subunits.

### Functional cell-type classification

Ganglion cells from marmoset recordings were classified based on their temporal filters and effective receptive field diameters. For each cell, the filter and diameter values were concatenated into a vector and then standardized to zero mean and unit variance across all cells. The vectors’ first few principal components that explained 0.9 of the variance were clustered using K-means clustering. The cells grouped by the cluster labels were checked by visual inspection, and occasional non-matching cells that violated receptive field tiling and temporal filter shape were removed from the cell cluster. The clusters were manually labeled as ON and OFF parasol and ON and OFF midget cells based on the sign and shape of the temporal filter and the diameter of the receptive field. Cells in an additional fifth cluster were considered unclassified and excluded from the analysis (excluded cells for each of the three recordings: 12 out of 109 cells total, 32 out of 127, and 39 out of 103).

### Spike-triggered stimulus ensemble

The spatial region of the stimulus relevant to a given ganglion cell’s response was defined as the smallest rectangular window, *x* × *y*, that encloses the three-sigma outline of the Gaussian fit of the receptive field (see above). The set of stimulus sequences that lie within this spatial window and within the 700 ms time window preceding a spike compose the spike-triggered stimulus ensemble. Frames in each spike-triggered stimulus were weighted by the temporal filter of the STA and then summed to one effective stimulus frame per spike. For multiple spikes occurring in one time bin, the identical effective stimulus was repeated accordingly. The effective stimuli of all spikes compose the effective spike-triggered stimulus ensemble (STE). The stimulus frames of the STE with *n* = *x* × *y* pixels were flattened into column vectors to form the STE matrix with dimensions *n* × *m*, corresponding to *n* pixels by *m* spikes.

### Sparse semi-non-negative matrix factorization

Semi-non-negative matrix factorization (semi-NMF) decomposes a matrix V ∈ ℝ*^n^*^×*m*^ into two factor matrices W ∈ ℝ*^n^*^×*r*^ and H ∈ ℝ*^r^*^×*m*^ with *r* < *m*, *n*, such that

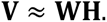

These variable names are common to NMF terminology and are related to STNMF in the following way: **V** is the STE matrix, where each column *v_i_* ∈ ℝ*^n^* is the effective stimulus frame of spike *i* ∈ 1, … , *m*. The columns of **W**, *w_k_* ∈ ℝ*^n^*, are the *k* ∈ 1, … , *r* spatial modules, and each column of **H,** ℎ*_i_* ∈ ℝ*^r^*, contains the corresponding *r* weights for each spike *i* ∈ 1, … , *m*. The stimulus *v_i_* that corresponds to spike *i* is approximated by the linear combination of the modules **W**, as basis vectors, and the corresponding weights, ℎ*_i_* , as coefficients or encodings. The product of **W** and **H** forms a lower-dimensional representation of **V** based on recurring structure within **V**. This gives rise to the basis spatial modules **W,** among which we find the resulting subunits.

Semi-NMF is an NMF variant that relaxes the non-negativity constraint on **V** and **H**, while **W** remains non-negative. To retain parts-based factorizations in semi-NMF, the loosened non-negativity constraint has to be replaced (Ding et al. 2010). Typically this is achieved by sparsity regularization on **W** (Hoyer 2002, 2004; Kim and Park 2007, 2008). Here, this resulted in the use of sparse semi-NMF.

NMF implementations generally improve the approximation **WH** iteratively by minimizing an objective function. As is common for NMF, we chose the Euclidean distance between **V** and the reconstruction **WH**, because it is simple to differentiate (Lee and Seung 1999). Sparsity was implemented with an *ℓ*1-norm penalty on the modules as an additional term in the objective function, weighted by a regularization parameter λ

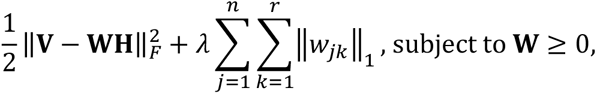

where w*_jk_* for *j* = 1, … , *n* and *k* = 1, … , *r* are the elements of **W**, ‖·‖^2^ is the squared Frobenius norm and ‖·‖_1_ is the *ℓ*1-norm.

To distinguish the localized subunits from noisy modules in **W**, the spatial autocorrelation was calculated using Moran’s I (Li et al. 2007; Moran 1950). Given module w*_k_* for *k* = 1, … , *r* as the vector *s* ∈ ℝ*^n^*, the autocorrelation was computed with

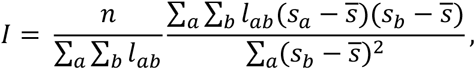

where *a*, *b* = 1, … , *n* are pairs of pixel indices, ̄*s* is the mean of vector *s*, and L ∈ ℝ*^n^*^×*n*^ is the neighbor matrix with *l_ab_* equal to unity if *a* and *b* correspond to spatially adjacent pixels in the two-dimensional layout of the *n* = *x* × *y* pixels and zero otherwise. The scalar *I* ranges from negative to positive unity with higher positive values denoting denser localization in space. We chose an autocorrelation threshold of *I* = 0.25 to distinguish localized subunits from noise modules. Additionally, one can consult the mean spike weight of a given module *k* by averaging the *k*th row of **H** to exclude candidate subunits from among the modules when their contribution to the spike generation is small, e.g., smaller than that of non-localized modules. We encountered such low-weight localized modules only rarely and excluded them from the final set of displayed subunits in a few cases based on this criterion.

### Accelerated fast hierarchical alternating least squares

The objective function was minimized by alternating optimization of the two factor matrices. Given a prior initialization of **W**, **H** was updated while **W** was held fixed, and **W** was subsequently updated while the newly computed **H** was held fixed. This process is generally repeated until a termination criterion is reached. We here chose a fixed number of 1,000 iterations (occasionally 200 or 5,000 for selected checks) as most cells showed convergence much earlier. The iterations concluded with a final update of **H** to provide the appropriate weights for the final set of modules. As the update of **H** is unconstrained, unlike the update of **W**, this also helps in obtaining an accurate assessment of the achieved reconstructions error.

The alternating updates of **W** and **H** split the optimization into two sub-problems. In the context of semi-NMF, **H** is not constrained to be non-negative. Given the Euclidean distance objective function, the update of **H** reduces to a least squares problem (Ding et al. 2010) and can be solved accurately using the Moore-Penrose inverse or pseudoinverse (·)^†^

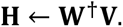

Subsequently, we applied an *ℓ*2-normalization of the rows of **H**. This normalization step is generally performed in Euclidean-distance-based NMF to counteract non-uniqueness of solutions (since a scaling of a row in **H** can be compensated by an opposite scaling of the corresponding column in **W** without affecting the reconstruction) and to prevent exploding coefficients during the alternating updates of **W** and **H** (Ho 2008). The update of the non-negative matrix **W**, on the other hand, poses a non-negative least squares (NNLS) problem, in fact a regularized NNLS in the case of sparse semi-NMF. HALS algorithms (Cichocki et al. 2007) approximate it by updating the modules sequentially. We incorporated our sparsity constraint into Fast HALS (Cichocki and Phan 2009) with the local update for *w_k_* for *k* ∈ 1, … , *r*

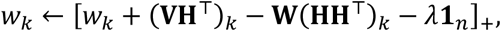

where *w_k_* and (·)*_k_* are the *k*th column of **W** and of the result of the matrix multiplication in parentheses, respectively, 1*_n_* ∈ ℝ*^n^* is a column vector of length *n* containing all ones, and [·]_+_ = max{·, 0} is the elementwise maximum with zero of a vector. The terms VH^T^ and HH^T^ remain fixed, whereas **W** is updated in-between the column updates. This update rule is based on Fast HALS with additional sparsity regularization. Fast HALS simplifies the update equation of HALS and results in a significant speed up (Cichocki and Phan 2009). It allows terms to be dropped from the HALS equation facilitated by the rows of **H** being *ℓ*2-normalized prior to the update. Fast HALS takes advantage of this normalization step.

While Fast HALS improves the local update rule on the scale of one module, A-HALS (Gillis and Glineur 2012) optimizes the computational cost on the scale of all modules in **W**. While **W** is updated continuously, A-HALS exploits the fact that the terms VH^T^ and HH^T^ in the local update remain unchanged over the course of iterating over the modules. These matrix multiplications outweigh the other terms of the local update rule in the number of floating-point operations, making the update of **W** costly compared to the outcome. A-HALS wraps an outer loop around the local updates of each column of **W**, effectively updating **W** multiple times per one NMF iteration, in order to tradeoff the floating-point operations in the pre-computed matrix multiplications and in the local updates. This increases the approximation at comparably little additional cost and improves the accuracy of a single update of **W**.

To determine the number of outer loop cycles, we found the previously suggested parameters to be suitable (Gillis and Glineur 2012). The maximum number of outer loop cycles is determined by *α* × *ρ*_W_, where 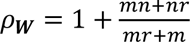 is the number of floating-point operations in the update and *α* = 0.5 is a scaling factor. An additional dynamic stopping criterion checks whether the last improvement of **W** is smaller than 10% of the first improvement, that is, in the first cycle of the outer loop. The improvement of **W** is measured as the Frobenius norm of the difference to the previous cycle of the outer loop, ||**W** – **W**^(*previous*)^||*_F_*. Aside from the acceleration by performing multiple update cycles, A-HALS improves numerical stability by ensuring no zero-columns in **W** to avoid convergence issues of HALS (Gillis and Glineur 2008; Ho 2008). Columns of all zeros are replaced by columns whose entries all take a small positive constant (here 10^−16^). In the context of semi-NMF, this facilitates, in particular, the computation of an SVD-based pseudoinverse W^†^ for the subsequent update of **H**. We combined Fast HALS and A-HALS by replacing the local update in A-HALS with the equations from Fast HALS and complementing this with sparsity regularization. All analyses were carried out on computers with an Intel Xeon CPU E3-1270 v5 @ 3.60 GHz.

### Non-negative singular value decomposition with low-rank correction

To obtain suitable, guided starting points for the alternating optimization of **H** and **W**, we initialized the modules **W** using non-negative singular value decomposition with low-rank correction (NNSVD-LRC), which had been introduced for full NMF (Atif et al. 2019). Here, we describe a modified version for semi-NMF with slight optimizations. Given *r* modules, we extracted half that number of components from a truncated SVD of **V**. Specifically, we used 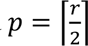 components from the decomposition V ≈ U**Σ**A with the matrices U ∈ ℝ*^n^*^×*p*^ and A ∈ ℝ*^p^*^×*m*^ and the diagonal matrix Σ ∈ ℝ*^p^*^×*p*^. To obtain a two-factor form as in NMF, the singular values were integrated into the left and right singular vectors with their element-wise square root to provide 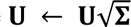 and 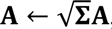, respectively, so that V ≈ UA with the modified U and A. Unlike the original NNSVD-LRC, we only initialized **W** (via U) and thus discarded A. The components of **U** sorted by decreasing singular values served as the columns of **W**. In doing so, each component was used twice, with and without a sign-flip, ordered so that the one with the largest positive element came before its sign-inverted counterpart to ensure reproducible component ordering. (In case of an odd *r*, the component with the lowest singular value was only added once.) In the end, all negative entries were set to zero to make the initialization of **W** non-negative. This resulted in the initialization **W** = [*u*_1_, −*u*_1_, *u*_2_, −*u*_2_, … , *u_p_*, −*u_p_*]_+_. Inspired by A-HALS, we implemented an additional step to avoid numerical instability (Gillis and Glineur 2012). Any columns of **W** that contained only zeros were set to a small positive value (10^−16^). Any zeros residing in non-zero columns remained to accelerate matrix multiplications and to allow sparsity to emerge.

The solution can be improved further at low computational cost by a few NMF iterations on the low-rank approximation V^(*p*)^ ≈ UA . As V^(*p*)^ has a factorization with low-rank matrices, the matrix multiplications for the NMF iterations become less complex, and NMF iterations reduce from a complexity of 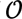(*nmr*) to 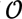((*n* + *m*)*r*^2^) (Atif et al. 2019; Gillis and Glineur 2012). In the original NNSVD-LRC, the low-rank NMF was performed using A-HALS until the difference in relative reconstruction error 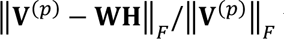 to the previous iteration fell below 5% of the initial error, which typically occurred within ten iterations (Atif et al. 2019). Here, since we used sparse semi-NMF instead, we observed that the proposed termination criterion was always reached within one AF-HALS update in the case of STNMF. We thus fixed the number of iterations with V^(*p*)^ at one and omitted the termination criterion. Without the need for computing reconstruction errors, it suffices to initialize **W** from the SVD components, leaving **H** to be subsequently computed from the initialization of **W**. This further reduces computational cost and provides a fast initialization procedure.

For comparison and for consensus methods, we applied random initialization of **W** by sampling its values independently from a uniform distribution between zero and unity. We used the Mersenne Twister (MT19937) pseudorandom number generator (Matsumoto and Nishimura 1998) for potential compatibility across Python, R, and MATLAB. For reproducibility, we applied a specified set of seeds, here ranging from 0 to 49 for the 50 repetitions of the consensus analysis.

### Consensus analysis

We used consensus analyses (Brunet et al. 2004; Monti et al. 2003) to determine the best tradeoff between solution stability and sparsity. Consensus methods allow judging the robustness of the sparsity parameter instead of assessing the reconstruction error, or residual. For NMF, the latter has been known to be problematic (Bro et al. 2008; Owen and Perry 2009) because leaving out samples from the training data requires fitting the weights corresponding to the samples on the test data, which biases the reconstruction error. Leaving out individual entries from the data matrix instead of full samples, on the other hand, could alleviate the problem of refitting weights, but are not compatible with the accelerated NMF algorithms applied here. Furthermore, in the context of STNMF, the noisy excess modules may influence the residual, but neglecting them instead, favors solutions that exhibit more localized modules over those with fewer. Consequently, decompositions based on different parameters cannot be compared easily by their residual. For these reasons, cross-validation is not suitable for STNMF, suggesting the application of consensus methods.

For applying the consensus analysis, we performed 30 randomly initialized repetitions of sparse semi-NMF (50 repetitions for the detailed comparative analysis) for a range of parameters. The parameters correspond to different sparsity regularization strengths. Semi-NMF was run for 1,000 iterations, which we found to typically be enough to reach convergence. Similarity between solutions was assessed by treating the encodings in **H** as cluster labels and comparing them across solutions. We define cluster labels as the index of the module with the highest absolute weight for a given spike. Although a spike may be elicited by a combination of subunit activations, considering the maximum associated module suffices for sake of comparison. Because the ordering of the emerging modules is arbitrary, the cluster labels cannot be compared across solutions directly. To remove this ambiguity, pairwise comparisons of cluster labels among the spikes within each solution are computed in a connectivity matrix **G**. Each entry *g_ij_* in the square matrix is unity, if spikes *i* and *j* share the cluster label and zero otherwise. As the number of unique pairs grows according to *m*(*m* − 1)/2 for increasing number of spikes *m*, we limited ourselves to 25000 spikes selected at random with a persistent selection across the analysis. That amounts to more than one gigabyte of memory at 32-bit floating-point precision for one decomposition. The number may be increased with more memory available or with memory-efficient implementations. Nevertheless, we found that this subsample provides a sufficient representation of the resulting cluster memberships. The elementwise average of the connectivity matrices of all 30 (or 50) solutions expresses the consensus across solutions. This consensus matrix contains values between zero and unity, describing the fraction of agreement between the encodings across the 30 (or 50) runs, and the cophenetic correlation coefficient (CPCC) quantifies how well these agreement values separate the data points (spikes) into groups, which here correspond to activation of individual subunits. The CPCC takes values between zero and unity, with unity denoting perfect consensus and identical solutions (Brunet et al. 2004). We calculated the cophenetic correlation by converting the connectivity matrix into a distance matrix with entries *d_ij_* (that is, by subtracting each entry from unity) and using (from the Python module “scipy.cluster.hierarchy”) the function “linkage” (with the “average” method) to perform hierarchical clustering, followed by the function “cophenet” to obtain the cophenetic distances within the clustering. The hierarchical clustering yields a dendrogram for the hierarchical joining of data points, with the cophenetic distance of two data points given by the height of the dendrogram (i.e., distance of joined clusters) when these data points are joined. The CPCC is then defined as the Pearson correlation between the original distances *d_ij_* and the cophenetic distances *c_ij_* across all pairs of data points:

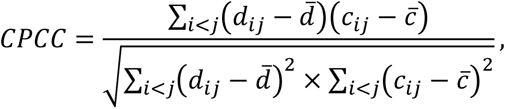

where ̄*d* and ̄*c* are the corresponding average distances.

STNMF reserves excess modules to capture noise in the data. These modules are not considered localized subunits and are not representative of the solution. As uncorrelated noise is distributed arbitrarily across the excess modules, they are likely to vary across solutions. Thereby they skew the described metric. Consequently, we only considered modules containing localized subunits as valid cluster labels. Connectivity pairs involving non-localized modules were set to zero in the connectivity matrices. These entries do not reach consensus across solutions. This causes a lower dispersion in the consensus matrix, quenching the cophenetic correlation coefficients, and the metric becomes more sensitive to differences in detected subunits. In the case of no localized subunits in the decomposition across runs, we did not compute the cophenetic correlation. This results in gaps in the stability curve (see **Figure 11A** for examples).

The cophenetic correlation coefficient corresponds to the similarity of the recovered subunits across the repeated decompositions. It represents the stability of the solutions, and we used it to compare the different sparsity regularization parameters. The optimal trade-off between high stability and little regularization occurs at the bend of the curve of regularization strength versus stability (**Figure 7C**). Here, we determined the inflection point by visual inspection and defined it as the final sparsity parameter. For the detailed population analyses, we therefore here used the following values for the sparsity parameter: 1.74 for salamander fast-OFF cells, 1.5 for marmoset ON parasol cells, 1.9 for marmoset OFF parasol cells, 2.0 for marmoset ON midget cells, and 2.5 for marmoset OFF midget cells. To instead identify the sparsity parameter value more systematically, we propose that one could describe the curve as a piecewise linear function with two or three components and select the intersection of the functions.

### Subunit comparison

To compare recovered subunits on a geometrical basis, we computed their relative spatial overlap. Given the contour outlines (described above) of two subunits *i* and *j*, we calculated their relative overlap *o_ij_* with the Jaccard index (Jaccard 1912), which is defined as the ratio of the intersection to the union of their areas *a_i_* and *a_j_*:

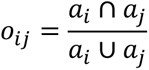

We considered subunit pairs with *o_ij_* > 0.5 , that is, more than 50% area overlap, to be strongly overlapping and thus candidates for identical subunits.

For the purpose of determining prevalent subunits across multiple randomly initialized runs of STNMF (**Figure 6**), the relative overlap was compared across decompositions. A subunit that showed overlap with *o_ij_* > 0.5 across more than 50% of the runs (colored outlines in **Figure 6**) was considered a robustly recovered subunit.

### Inter-mosaic coordination

The spatial coordination of ON- and OFF-type mosaics (both for ganglion cell receptive fields and subunits) was determined using the inter-mosaic coordination energy (IMCE) as proposed previously (Roy et al. 2021). In brief, individual receptive fields or subunits were reduced to centroids by their center of mass. Within a region of interest, one of the mosaics was shifted in horizontal and vertical position relative to the other. The squared inverse distance between each heterotypic pair of centroids *q_ij_* was measured with

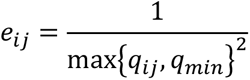

where *q_min_* is a minimum distance set to 0.2 times the median heterotypic nearest-neighbor distance. The values of *e_ij_* were averaged across all pairs to obtain the IMCE. The IMCE was computed for different offsets of the OFF-type mosaic relative to the ON-type mosaic on a 50-by-50 Cartesian grid before being z-scored. The shift distances were normalized to the median homotypic nearest neighbor distance of the ON-type (frozen) mosaic. The generated topographic map of IMCE values was radially averaged to examine whether the IMCE increased or decreased with radial shift distance. We selected the analyzed region of interest in two ways. First, we manually chose a region where the density of subunits was approximately homogeneous on both mosaics to minimize effects of gaps. We then confirmed the determined IMCE by following the proposed implementation of finding a suitable convex hull that maximized the number of enclosed heterotypic pairs while excluding outliers (Roy et al. 2021).

